# Macrophage Phagocytic Impairment is Associated with Dysbiosis of the Respiratory Microbiome in Frail Older Adults

**DOI:** 10.64898/2026.02.09.703495

**Authors:** KBR Belchamber, KP Yip, OS Thein, FS Grudzinska, S Patel, E Sapey, T Jackson, D Parekh, MJ Cox, A Scott

## Abstract

Advanced age and frailty are risk factors for pulmonary infections, which are the leading cause of death in older adults. The mechanism behind increased susceptibility to infection is poorly understood, but interactions between innate immunity and the respiratory microbiome may be implicated. We investigated changes to monocyte-derived macrophage function, and the respiratory microbiome in healthy young adults, healthy older adults, and frail older adults. We found that macrophage phagocytosis and efferocytosis is impaired in frail older adults, associated with elevated pro-inflammatory cytokine secretion and a failure to regulate expression of phagocytic receptors CD14 and CD36. Pre-treatment of macrophages with a PPARγ agonist restored phagocytosis, implicating this pathway in cellular dysfunction. Accompanying respiratory microbiome showed reduced bacterial diversity and richness in older adults, which was more advanced in frail older adults. These results are consistent with a macrophage predation defect in the lungs during frailty which may impact bacterial diversity, and may be improved via manipulation of the PPARγ pathway.

**Graphical abstract:** 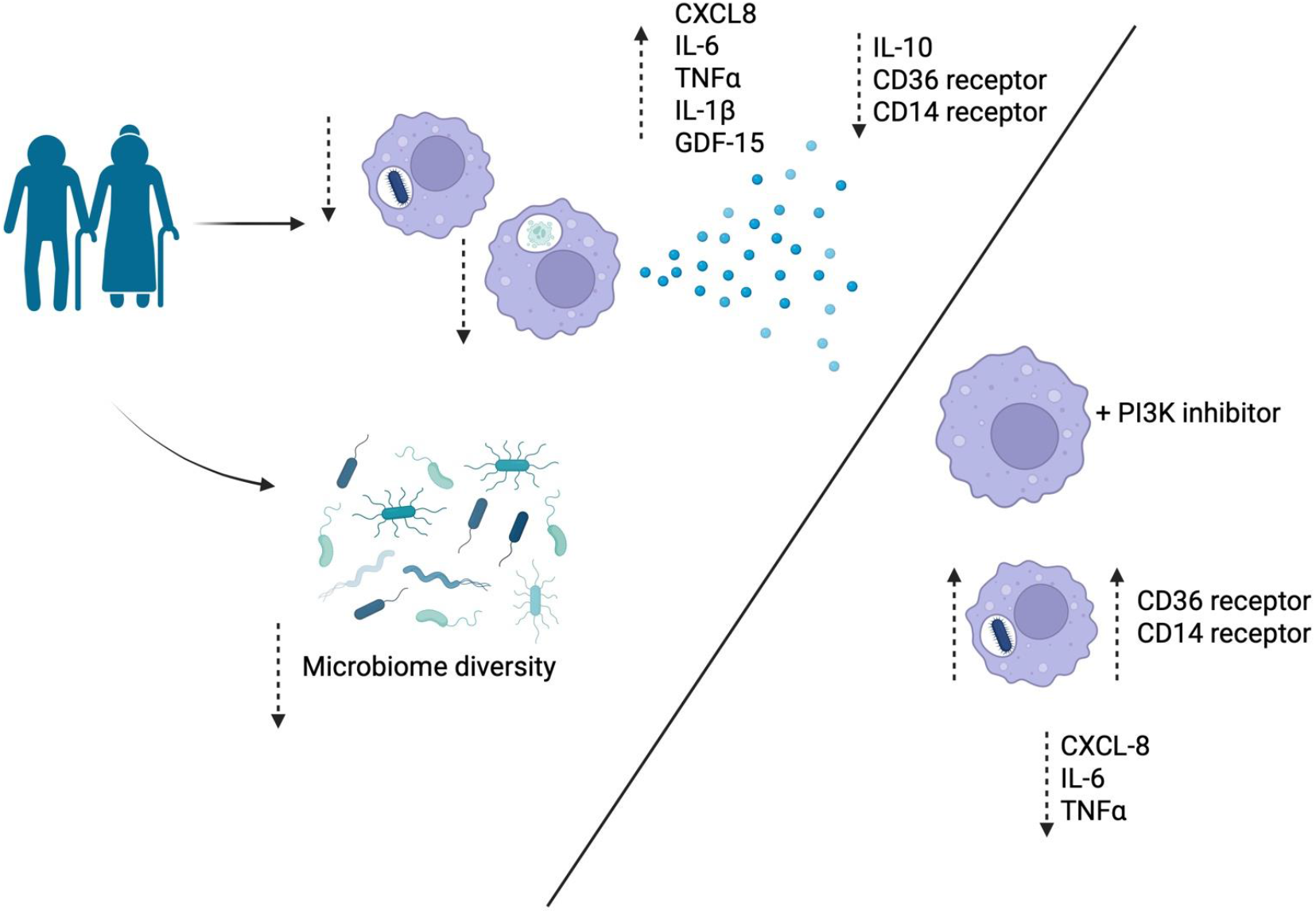

## Background

Advanced age is a predominant risk factor for pulmonary infections, which are the leading cause of death in older adults. Over 70% of pneumonia cases requiring hospital treatment occur in people aged over 65, with 20% mortality in this age group ^1^. ‘Inflamm-ageing’ is thought to be central to this process^2^, characterised by increased circulating and tissue pro-inflammatory cytokines, including C-reactive protein (CRP), Interleukin (IL)-6, and Tumour necrosis factor (TNF)*α* ^3, 4^, linked to an increased susceptibility to chronic morbidity, disability and frailty ^5, 6^. Ageing is also linked to declining health overall, with increased age associated with higher incidence of co-morbidities including frailty. Frailty is defined as a gradual loss of the body’s reserves, making an individual more vulnerable to sudden health changes and is an independent predictor of poor outcomes and heightened susceptibility to infection ^7, 8^.

The innate immune system is heavily implicated in the pathophysiology of age-related lung diseases including respiratory infections^9^, with impaired neutrophil function described in older adults with pneumonia which does not recover in survivors^10^. In mice, advanced age is linked to impaired alveolar macrophage (AMφ) phagocytosis, antiviral responses, and Toll like receptor (TLR) signalling ^11^, as well as impaired bacterial clearance, pro-inflammatory cytokine release and features of mitochondrial dysfunction ^12^. Aged murine AMφ display decreased ATP production, enhanced reactive oxygen species (ROS) generation, and diminished antioxidant responses to respiratory bacteria ^12^. Due to species differences in myeloid cells between mice and humans ^13^, these studies require validation in humans, however they do suggest that impairment in macrophage function in the lungs may contribute to lung damage and to the decline in lung function observed in normal ageing ^14^. In humans, impaired macrophage function, characterised by reduced phagocytosis and bacterial killing, and altered inflammatory mediator release, is linked to age related diseases including chronic respiratory disease (COPD, IPF) and cancer ^15^, but how macrophage function relates to healthy ageing is unclear.

Studies attempting to identify clinical factors associated with impaired innate immune cell function in ageing have discovered that frailty is more important than ageing alone ^16^. Frail older adults exhibit increased systemic inflammation ^17^, altered adaptive immune cell populations ^18^, and impaired neutrophil function ^19^. The causes and mechanisms underpinning these changes are not known, but micro-environmental factors are likely sources. These range from the presence of systemic or local inflammation^20^ to changes in the local environment including the microbiome^21^.

The upper respiratory tract microbiome reveals reduced bacterial density and reduced diversity with age ^22, 23^, with the changes associated with frailty understudied. One important study showed a loss of species richness associated with increased frailty in older adults, but in a cohort of hospitalised patients that received peri-operative antibiotics, which may have skewed results ^24^. Pneumonia is further associated with a reduced microbial diversity in the respiratory tract and often caused by bacteria that can either be acquired or carried in health, such as *Streptococcus pneumoniae*, and *Haemophilus influenzae*^25^.

The association between altered immune cell function in ageing and age-related disease, and changes to the respiratory microbiome seen with age require investigation. It is unclear whether impaired immune cell function enables to respiratory microbiome to become permissive to infections by pathobionts (bacteria which normally colonise humans, but have potential to become pathogenic). Further, it is unclear whether in frailty this interaction is increasingly impaired, leading to an increased risk of infection and poorer outcomes. Understanding these interactions may give us tools to reduce the morbidity and mortality associated with increased infection risk with increasing age. In this study we hypothesised that human monocyte-derived macrophage function declines with age and frailty, and that this is associated with altered respiratory microbiome.

## Methods

### Participants

All participants were recruited in accordance with the HRA and ethically approved protocol (18_WM_0097: West Midlands Solihull). Inclusion and exclusion criteria for each group are provided in table 1. Frail older adults (age 65-100) were recruited as inpatients at the Queen Elizabeth Hospital Birmingham, admitted due to social or mild clinical need, but were assessed as being free from infection or bodily trauma, and categorised as frail according to the Clinical Frailty Scale^26^ by admitting physician. Non-frail older adults (age 65-80) were recruited from volunteers from the 1000 elders cohort^27^, categorised as non-frail according to the Clinical Frailty Scale, with the absence of any acute medical conditions. Healthy young adults (age 21-35) were recruited from staff members at the University of Birmingham Research Laboratories in the Queen Elizabeth Hospital, Birmingham.

**Table 1.** Cell surface markers antibodies used to define key monocyte populations.

| Marker | Fluorophore | Cell type | Volume | Cat number |
| --- | --- | --- | --- | --- |
| CD14 | Vioblue | Monocyte | 5ul | Biolegend<br>367144 |
| CD16 | APC-Vio770-REA | Monocyte | 2ul | 130-113-390 |
| CD3 | Viogreen-REA | T cells | 2ul | 130-113-142 |
| CD56 | PE-Vio615-REA | NK cells | 2ul | 130-114-550 |
| CD19 | Viobright-515-REA | B Cells | 2ul | 130-113-650 |
| CCR2 | APC-REA | Monocyte activation | 2ul | 130-109-596 |
| CCR5 | PE-REA | Monocyte activation | 2ul | 130-117-356 |

**Table 1.** inclusion/exclusion criteria.

| Inclusion | Exclusion |
| --- | --- |
| Age between 18-35 years (young adults) | History of respiratory disease, or significant respiratory symptoms including breathlessness and cough |
| Age between 65-100 years (older adults) | Hematological malignancy |
| Frailty score between 1-8 | Infection or antibiotics use in previous 2 weeks |
| Provision of informed written consent | Immunosuppressive medications |

### Sample collection

Participants underwent venepuncture to collect fresh venous blood into lithium-heparin vacutainers for cellular experiments and plasma collection, or silica vacutainers for serum collection. Serum tubes were allowed to clot for 30 minutes at room temperature, and then both serum and plasma tubes centrifuged at 500 g for 10 minutes. Plasma or serum was removed and stored in cryovials at −80C. Participants also underwent an oropharyngeal swab of the posterior oropharyngeal membrane on both sides, for 5 seconds using dry cotton swabs (Copan) according to established protocols^28^. Swabs were immediately transferred to and stored at −80C.

### Monocyte isolation

20 mL of blood collected in lithium heparin was used for monocyte isolation by Percol® gradient centrifugation^29^, with the PBMC layer collected and transferred to a fresh 15ml falcon tube. Cells were washed by centrifugation in PBS at 300 g for 5 minutes, then resuspended in RPMI (Gibco) + P/S + L-Glutamine. Monocytes were counted by haemocytometer and diluted to 1×10^6^ cells/ml for experimentation. Experiments were prioritised based on cell count, which is reflected in the number of subjects for each experiment.

### Monocyte phenotyping

1×10^6^ PBMC were transferred to a FACS tube, washed in PBS (300 g, 5 mins) and resuspended in FACS buffer. Cells were blocked with 10% human serum for 10 minutes, then washed with FACS buffer. 100 μl of cell suspensions were subsequently stained with mouse anti-human monoclonal antibodies including fluorescence minus one (FMO) controls, for 10 minutes at 4°C, washed and resuspended in FACS buffer. Table 1 shows antibodies used. Flow cytometry was performed on a MACSQuant10 (Surrey, UK), and analysed using Flowjo software, with 100,000 PBMC recorded for each sample. Gating strategy can be seen in Supplementary Figure 1.

### Monocyte-derived macrophages

Monocytes were plated in 24-well plates at 0.5×10^6^ cells/well and left to adhere for 2 hours. Media was removed by aspiration and replaced with fresh media containing 2 ng/ml GM-CSF. Media was replaced every 3 days, until day 10 when cells were confirmed as MDM by microscopy.

### Bacteria

*Streptococcus pneumoniae* (Serotype 14, NCTC 11902, National Collection of Type Cultures) was grown as previously described [27]. Non-typeable *Haemophilus influenzae* (NCTC 1269, National Collection of Type Cultures) was cultured on chocolate agar overnight and then grown to an optical density at 600 nm of 0.6 in Brain Heart Infusion broth (cat. no. CM1135, Oxoid Ltd, Basingstoke, UK) supplemented with 10 μg/mL heme (cat. no. H9039, Sigma-Aldrich Company Ltd). Bacteria were not opsonised. Heat-killed bacteria were generated by incubation at 65°C for 2 hours as described previously [25].

### Fluorescent labelling of heated-killed bacteria

Bacterial cultures were fluorescently labelled using AlexaFluor 404 NHS ester (cat. no. A30000, Life Technologies, Paisley, UK) or AlexaFluor 488 NHS ester (1 mg·mL−1 in dimethyl sulfoxide (DMSO), cat. no. A20100, Life Technologies) and incubated overnight. Labelled bacteria were washed repeatedly in PBS to remove unbound label, then resuspended in PBS and stored at −20°C.

### Generation of apoptotic neutrophils

Neutrophils were isolated from peripheral blood of healthy young donors as previously described ^29^. Neutrophils were stained with PKH-26 red fluorescent dye (Sigma) according to manufacturer’s instructions. Staining was stopped by addition of 1% (w/v) BSA. Cells were in RPMI before being resuspended at 5.0 x 10^6^ cells/ml in RPMI supplemented with 10% (v/v) FCS. Cells were then cultured for 20 h. This resulted in >80% apoptotic and fewer than 5% necrotic cells as measured by Annexin V staining (data not shown).

### Bacterial phagocytosis and efferocytosis

For phagocytosis, fluorescent bacterial stocks were sonicated then added to macrophages at a ratio of 80:1 and incubated at 37 °C for 4 h. For efferocytosis, apoptotic neutrophils were added to macrophages at an MOI of 4 and incubated at 37 °C for 4 h. Media was then removed and stored at −80C for cytokine analysis. Cells were then washed twice with PBS to remove free bacteria/neutrophils and dissociated from the plate using cell dissociation media and vigorous pipetting. Cells were transferred to FACS tubes and fixed with 4% PFA for 10 minutes at room temperature. For rosiglitazone treatment, frail older adult MDM were pre-treated with 1*μ*M rozigliazone or vehicle control (0.01% DMSO) for 16 hours, prior to phagocytosis assay. Results are expressed as % total macrophages that phagocytosed prey, or as median fluorescent intensity (MFI), which indicates the amount of prey phagocytosed, expressed as mean ± standard deviation. Gating strategy shown in Supplementary Figure 2.

### MDM phenotyping

After phagocytosis/efferocytosis, cells were washed, then resuspended in FACS buffer, and incubated with 10% human serum for 10 minutes to block. Cells were washed and resuspended in FACS buffer for labelling with mouse anti-human monoclonal antibodies including fluorescence minus one (FMO) controls for 10 minutes at 4 °C. Cells were then washed and resuspended in FACS buffer. Table 2 shows antibodies used. Flow cytometry was performed on a MACSQuant10 (Surrey, UK), and analysed using Flowjo software, with 100,000 PBMC recorded for each sample. Alexa-488 labelled bacterial phagocytosis was measured in the FITC channel, whereas PHK26 labelled efferocytosis was measured in the PE channel (TLR2 antibody not included in this tube). Gating strategy shown in Supplementary Figure 2.

**Table 2.**
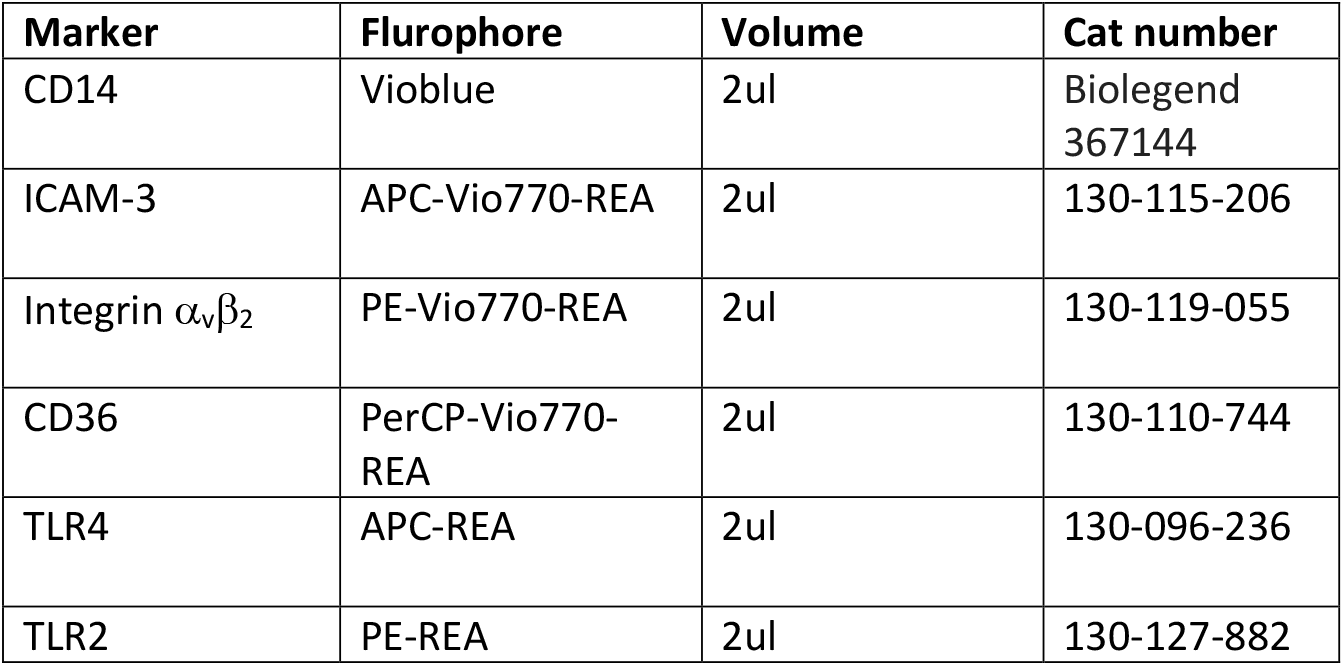
Cell surface marker antibodies used to investigate scavenger receptor expression.

**Table 2.** demographics.

|  | Young | Old | Frail |
| --- | --- | --- | --- |
| Male:Female | 7:5 | 3:9 | 5:7 |
| Age | 29 (21-35) | 75 (66-80)** | 82 (67-90)*** |
| White:non-white | 6:6 | 12:0 | 9:3 |
| Frailty score | 1 (1) | 2 (1-3) | 6 (4-8)***/### |
| Comorbidities |  |  |  |
| CVD | 0 | 3 (25%) | 9 (75%) |
| HYP | 0 | 4 (33%) | 7 (58%) |
| END | 0 | 0 (0%) | 3 (25%) |
| Arthritis | 0 | 1 (8%) | 4 (33%) |
| Other | 0 | 2 ( 16%) | 5 (41%) |

### Measurement of mROS using MitoSOX assay

Post phagocytosis, wells were washed with Hanks’ balanced salt solution (HBSS) (calcium free) and 5 μM MitoSOX dye (cat. no. M36008, Thermo Fisher, Loughborough, UK) added as per manufacturer’s instructions. Cells were incubated for 30 min at 37°C, then washed with HBSS containing calcium (0.185 g·L−1). Cells were dissociated from the plate using cell dissociation media and vigorous pipetting. Cells were transferred to FACS tubes, centrifuged, resuspended in 200 μL HBSS and analysed by flow cytometry using a MACSQuant 10 (Surrey, UK), in the B2 channel, and bacteria were measured in the V1 channel. Gating strategy can be found in Supplementary Figure 3.

### Measurement of mitochondrial membrane potential using JC-1

To measure mitochondrial membrane potential (MMP, ΔΨm), a tetrachloro-1,1’,3,3’-tetraethylbenzimida-zolylcarbocyanine iodide (JC-1) assay was performed post phagocytosis as per manufacturer’s instructions (cat. no. M43152, Thermo Fisher). Briefly, cells in a 24-well plate were washed with PBS and replaced with 500 μL warm cell culture media. 50 μM carbonyl cyanide m-chlorophenyl hydrazone (CCCP) was added to the positive control well and the plate was incubated for 5 min at 37°C. 2 μM JC-1 dye was then added to all wells for a further 25 min at 37°C, after which the cells were washed with PBS. Cells were dissociated from the plate using cell dissociation media and vigorous pipetting. Cells were transferred to FACS tubes, centrifuged, resuspended in 200 μL PBS and analysed by flow cytometry using a MACSQuant 10 (Surrey, UK), in the B2 and B1 channel, and bacteria were measured in the V1 channel. MMP was expressed as percentage green monomers – whereby increased monomers indicates decreased MMP. Gating strategy can be found in Supplementary Figure 3.

### MDM Cytokine analysis

MDM supernatants were analysed for cytokine release by custom Luminex assay according to manufacturer’s instructions. Release of IL-12/IL-23p40 was below the level of detection.

### Serum cytokine analysis

Serum cytokines were analysed by custom Luminex assay according to manufacturer’s instructions. Release of GM-CSF, IFNγ, IL-1*β*, IL-10, IL-12p70, IL-2, IL-4, IL-5, IL-6, IL-8 and TNF*α* were below the level of detection. Data are presented as median (IQR).

### Upper respiratory microbiome analysis

Throat swabs were taken from 20 frail older adults, 27 older adults and 13 younger adults enrolled in the study, with demographic data for this group seen in 3 text. DNA extraction of swab heads was performed using the ZymoBiomics DNA/RNA miniprep kit (Zymo Research) with bead-beating of the swab head using Lysing Matrix E tubes (MPBiomedicals). Spin baskets were used to maximise yield of crude extract from swab-heads ^30^. Extracted DNA was submitted to Novogene’s 16S rRNA gene sequencing service and the V3-V4 region sequenced using primers 341F and 806R. Raw data was processed using Mothur and analysed in the R statistical environment using the PhyloSeq package. Detailed microbiome analysis methods can be seen in Supplementary text. 16S rRNA gene sequence data has been submitted to the European Nucleotide Archive under bioproject ID PRJEB107714. A permutational multivariate ANOVA model was constructed using ADONIS^31^ using all available technical, immunological and demographic variables, minimizing the number of variables and maximizing the variance in beta diversity (between sample diversity) explained. To identify which OTUs (operational taxonomic units, deemed equivalent to “species”) were different between the subject groups DeSeq2 and ANCOM-BC2 were used and only OTUs significantly different by both methods were considered significant.

### Statistical analysis of immunology data

Statistical analyses were performed using GraphPad Prism (version 9.0.2 for Mac, Dotmatics, Boston, Mass). A normality test (Shapiro-Wilk test) was carried out on all data sets. One way ANOVA (for normally distributed data) and the Kruskal-Wallis/Friedman tests (for not normally distributed data) were used for appliable data. Two-way repeated measures ANOVA was used for applicable data. A P-value of less than or equal to 0.05 was considered statistically significant.

## Results

### Clinical characterisation

Cellular experiments were performed on samples taken from 36 participants in all, 12 in each group of frail older, non-frail older, and younger adults. Demographics and comorbidities are listed in table 2.

### Inflammatory changes in frail older adults

Serum markers associated with inflammageing and frailty including VEGF-A (527pg/ml (3358) vs. 262pg/ml (896) vs. 209pg/ml (814), p=0.0352), CRP (7.6pg/ml (14.5) vs. 0.9pg/ml (1.9) vs. 0.65pg/ml (1), p=0.0015) and TIMP-1 (712pg/ml (350) vs. 630 (296) vs. 562pg/ml (386), p=0.0045) were significantly elevated in frail older adults compared to both non-frail older, and healthy young adults, while GDF-15 was elevated in both frail older (629pg/ml (1248), p<0.0001) and non-frail older adults (237pg/ml (597) vs. 1(33), p=0.0065, 4Figure 1).

**Figure 1.**
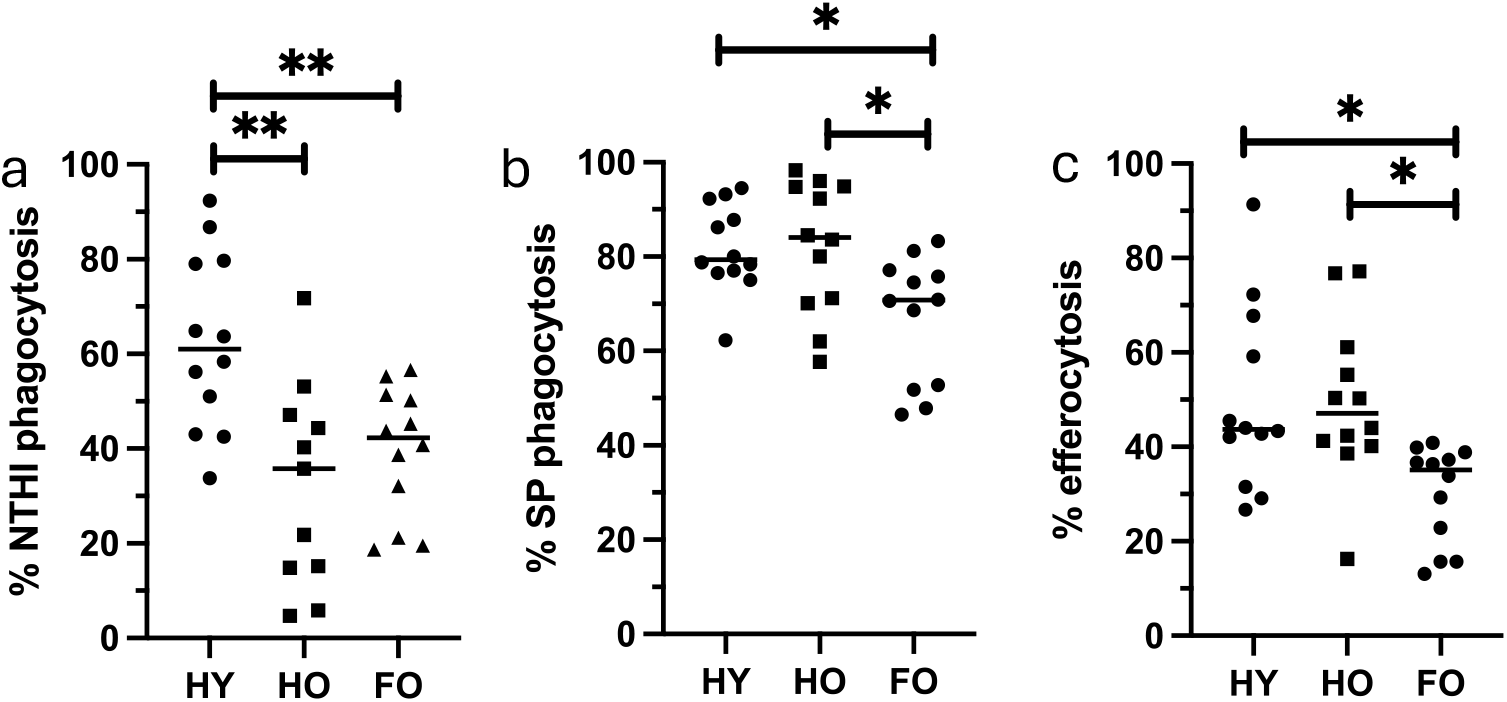
Monocyte derived macrophage function. (A) MDM phagocytosis of NTHI is significantly impaired in both HO adults, and FO adults compared to HY adults (HY 62±5% vs HO 32±6% vs FO 39 ±4%, p<0.001 and p<0.05). (B) MDM phagocytosis of SP is significantly impaired in FO adults compared to both HY and HO adults (HY 82±3% vs HO 82±4% vs FO 67±4%, p<0.05 and p<0.05). (C) MDM efferocytosis of apoptotic neutrophils is significantly impaired in FO adults compared to both HY and HO adults (HY 50±6% vs HO 50±5% vs FO 30±3%, p<0.05 and p<0.05). HY n=12, HO n=12, FO n=12. Analysed by One Way ANOVA.

Systemic monocyte populations were assessed via flow cytometry. Here, significantly reduced classical monocyte populations (CD14^++^ CD16^−^) were noted in frail older adults (HO 62% vs FO 76%, p<0.05, Supplementary Figure 5a), with elevated expression of CCR2 (p<0.01, Supplementary Figure 5b), a marker of monocyte recruitment from bone marrow^32^. There were also elevated intermediate monocyte populations (CD14^+^ CD16^+^) in frail older adults (HO 16.5% vs FO 11%, p<0.05, Supplementary Figure 5d), but similar levels of non-classical monocytes (CD14^−^ CD16^+^) were found (HO 7.5% vs FO 5%, p>0.05, Supplementary Figure 5g) with significantly reduced CCR2 expression in frail older adults (p<0.05, Supplementary Figure 5h), confirming previous reports^33, 34^.

### Macrophage function is impaired in frail older adults

To investigate macrophage function in frailty compared to non-frail ageing, we assessed MDM phagocytosis of respiratory bacteria. A significant reduction in the ability of non-frail older adult MDM to phagocytose *H. influenzae* compared to healthy young adults was seen (63± 19% vs. 40±13%, p=0.001, Figure 1a), but uptake of *S. pneumoniae* and efferocytosis remained comparable between non-frail older adults and healthy young adults. We also saw a significant reduction in the ability of frail older adult MDM to phagocytose *H. influenzae* (63± 19% vs. 32± 21%, p=0.0075, Figure 1a) compared to healthy young adults; and a reduction in the ability of frail older adult MDM to phagocytose *S. pneumoniae* (82± 9% vs. 82± 14% vs. 67± 13%, p=0.0139, Figure 1b) and to efferocytose apoptotic neutrophils (50± 19%vs. 50± 17% vs. 30± 10%, p=0.0155, Figure 1c) compared to non-frail older adults and healthy young adults. There was no change in MFI across the groups (Supplementary Figure 6).

MDM cytokine release was assessed after phagocytosis, to determine the effect on inflammatory responses. At baseline and compared to both healthy young and healthy older adult MDM, frail older adult MDM released increased levels of IL-6 (p<0.001 and p<0.001), TNFα (p<0.001 and p<0.001), IL-1β (p=0.0197 and p=0.0144), and GDF-15 (p=0.0196 and p=0.0033), suggesting that frail macrophages are pro-inflammatory at rest (Figure 2). There was no baseline difference in CXCL-8, IL-10, TIMP-1 or IFNγ between groups (p>0.05, Figure 2).

**Figure 2.**
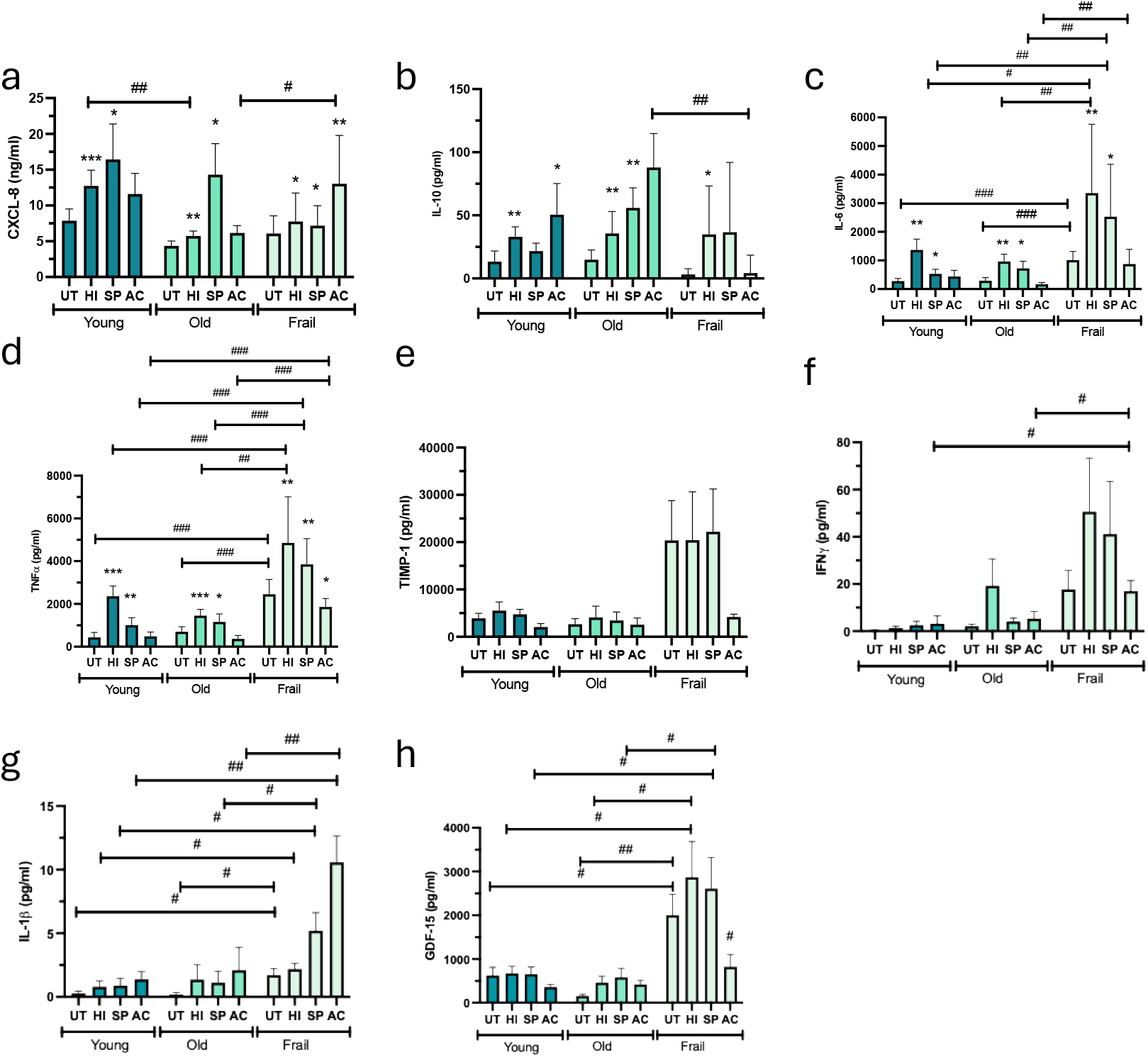
Cytokine release by MDM. (A) Phagocytosis of NTHI and SP induces increased release of CXCL-8 compared to UT cells, by all groups (p<0.05-p<0.001). Release of CXCL-8 by HO adults after NTHI phagocytosis was impaired compared to HY adults (12.8pg/ml vs. 5.7pg/ml, p<0.01). Release of CXCL-8 by FO adults after efferocytosis was significantly increased compared to HO adults (6.1pg/ml vs 13.0pg/ml, p<0.05). (B) Phagocytosis of NTHI and SP induces increased release of IL-10 compared to UT cells, by all groups (p>0.05, p<0.05-p<0.01). Efferocytosis induces increased IL-10 release by HY and HO adults, but not by FO adults (HY 50.1pg/ml vs HO 88pg/ml vs. FO 4pg/ml), which was significantly impaired (p<0.01). (C) Phagocytosis of NTHI and SP induces increased release of IL-6 compared to UT cells, by all groups (p<0.05-p<0.01). Baseline release of IL-6 was significantly increased in FO adults, compared to HY and HO adults (HY 279pg/ml vs. HO 291pg/ml vs FO 1012pg/ml, p<0.001). A similar pattern of increased IL-6 release by FO MDM was seen after phagocytosis of NTHI (p<0.05-p<0.01), SP (p<0.01) and efferocytosis (p<0.01). (D) Phagocytosis of NTHI and SP significantly increased release of TNFa compared UT cells by all groups (p<0.05-p<0.001). Baseline TNFa release was significantly increased in FO adults, compared t HY and HO adults (HY 435pg/ml vs. HO 699pg/ml vs FO 2455 pg/ml, p<0.001). Increased TNFa release by FO MDM was seen after phagocytpsos of NTHI (p<0.001) and SP (p<0.01), and efferocytosis (p<0.001). (E) TIMP-1 release did not change significantly across patient groups (p>0.05) or treatments (p>0.05. (F) Release of IFNg was significantly increased after efferocytosis in FO adults compared to HY and HO adults (HY 3.25pg/ml vs. HO 5.32 pg/ml vs. FO 17.02pg/ml, p<0.05). (G) Baseline release of IL-1B was significantly increased in FO adults compared to HY and HO (HY 0.27pg/ml vs HO 0.19pg/ml vs FO 1.7pg/ml, p<0.05). IL-1B release after phagocytosis of NTHI (p<0.05), and SP (p<0.05) and efferocytosis (p<0.01) was also significantly increased in FO adults compared to both HY and HO adults. (H) In FO adults, efferocytosis significantly impaired GDF-15 release compared to UT control (p<0.05). Baseline release of GDF-15 was significantly increased in FO adults compared to HY and HO (HY 634pg/ml vs HO 155pg/ml vs. FO 2001pg/ml, p<0.05). GDF-15 release after phaogocytosis of NTHI and SP was significantly increase in FO adults compared to HY and HO adults (p<0.05). (I) HY n=12, HO n=12, FO n=12. Analysedby Two-Way ANOVA.

Phagocytosis of bacteria by macrophages, including *H. influenzae* and *S. pneumoniae*, elevates macrophage release of pro-inflammatory cytokines as a response to infection^35^. We observed that frail older adult MDM released significantly increased levels of IL-6 (NTHI p=0.0277 and p=0.0086, SP p=0.0048 and p=0.0100, Figure 2c), TNFα (NTHI p=0.006 and p=0.0003, SP p=<0.0001 and p<0.0001, Figure 2d), IL-1*β* (NTHI p=0.0442, SP p=0.0137 and p=0.0258, Figure 2g) and GDF-15 (NTHI p=0.0231 and p=0.0146, SP p=0.0218 and p=0.0188, p<0.05, Figure 2h) after phagocytosis of either bacterium and compared to both non-frail older and healthy young macrophages, promoting inflammation locally.

Efferocytosis of apoptotic cells, in contrast, is an anti-inflammatory process which reduces pro-inflammatory cytokine release, and elevates anti-inflammatory cytokine release including IL-10^36^. Compared to baseline, both healthy young (p=0.0481) and non-frail older adult macrophages (p=0.0070) released IL-10 after efferocytosis with no change in pro-inflammatory cytokine release (Fig 2). In contrast, macrophages from frail older adults, did not increase IL-10 release after efferocytosis (p>0.05, Figure 2b), but released higher levels of CXCL-8 (p=0.0063, Figure 2a) and IL-1*β* (p=0033, Figure 2g) compared to baseline. This failure to initiate an anti-inflammatory response is likely detrimental to the resolution of inflammation, and supports the sustained inflammation seen in frail adults.

### Mitochondrial function in MDM

In order to determine whether changes in macrophage function could be attributed to mitochondrial function, mitochondrial ROS (mROS) and MMP were assessed. Compared to baseline, phagocytosis of *H. influenzae* and *S. pneumoniae* significantly increased mROS production by healthy young (p<0.001 and p=0.004), non-frail older (p<0.001 and p<0.001), and frail older MDM (p=0.0059 and p=0.0002), indicating a method by which MDM process bacteria. There were no differences in mROS production between subject groups. When MMP was assessed, the positive control CCCP significantly increased the percentage of green monomers across all subject groups (p<0.001) but there were no effects seen after phagocytosis, or across subject groups (Supplementary Figure 7).

### Macrophage scavenger receptor expression

Phagocytosis and efferocytosis are both processes mediated by scavenger receptors expressed on the macrophage surface, which bind their prey and initiate internalization. To assess whether altered expression of these receptors is responsible for impaired phagocytosis, we analysed expression of scavenger receptors on the surface of MDM at baseline, and alongside phagocytosis.

CD14, an important bacterial receptor^37^, was expressed by macrophages regardless of subject group or treatment, however frail older adult MDM expressed significantly reduced CD14 at baseline (74% vs. 47%, p=0.014), after phagocytosis of *H. influenzae* (84% vs. 60%, p=0.0149) and after efferocytosis (81% vs. 53%, p=0.0028) compared to healthy young MDM (Figure 3a). CD36, a receptor involved in both efferocytosis and phagocytosis^38^, was expressed consistently in healthy young macrophages regardless of treatment. However, in both healthy older and frail older macrophages, baseline expression of CD36 was significantly reduced (53% vs. 30% (p=0.0435) vs. 34% (p=0.033), Figure 3b). While expression of CD36 was elevated by these macrophages after phagocytosis of both *H. influenzae* (30-49%, p=0.009 and 34-50%, p=0.0154) and *S. pneumoniae* (30-59%, p=0.0079 and 34-56%, p=0.0349), it remained at levels significantly below that of healthy young macrophages (p=0.0081 and p=0.0369). After efferocytosis, both healthy older and frail older macrophages elevated CD36 expression to levels similar to that of healthy young macrophages (64% vs. 65% (p=0.0002) vs. 66% (p=0.0006), Figure 3b). Overall, impaired expression of, or recycling of receptors by older and frail older macrophages is implicated in impaired phagocytosis and efferocytosis. There were no significant differences in the % expression of ICAM-3, integrin, TLR2 or TLR4 across subject groups (Supplementary Figure 8), or in the MFI of all receptors analysed (Supplementary Figure 9).

**Figure 3.**
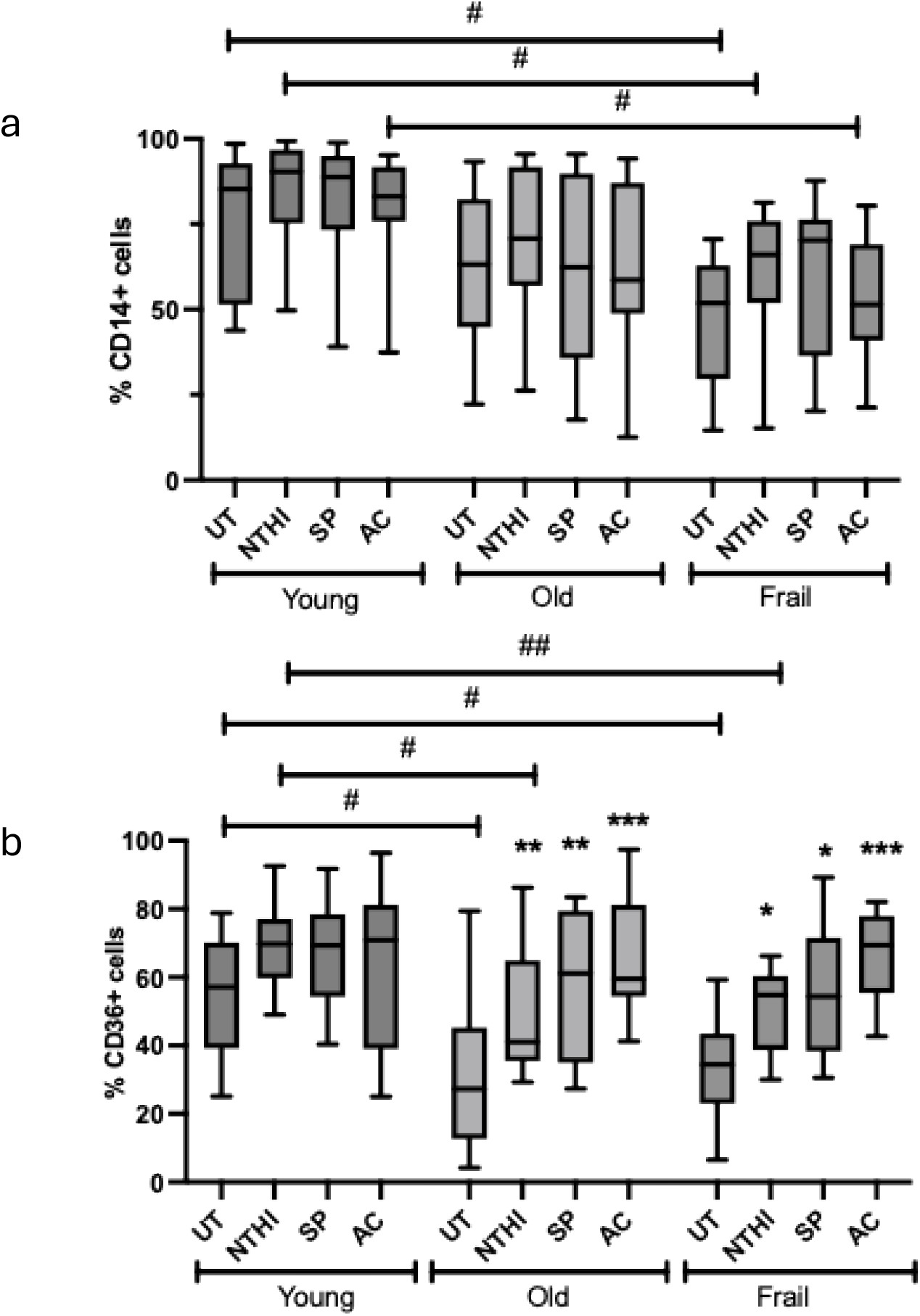
Cell surface receptor expression. (A) Compared to HY MDM, expression of CD14 was significantly reduced in FO MDM at baseline (74.3% vs 47%, p<0.05), after phagocytosis of NTHI (84% vs. 60%, p<0.05) and after efferocytosis (81% vs. 53%, p<0.05). (B) In HY MDM, there was no change in receptor expression across treatments. In HO MDM, phagocytosis of NTHI (p<0.01), SP (p<0.01) and efferocytosis (p<0.001) increased CD36 expression. Similarly, in FO MDM, phagocytosis of NTHI (p<0.05), SP (p<0.05) and efferocytosis (p<0.001) increased CD36 expression compared to baseline. Compared to HY MDM, expression of CD36 was significantly reduced in HO and FO MDM at baseline (HY 53% vs HO 30%, p<0.05, vs. FO 34% p<0.05), and after phagocytosis of NTHI (HY 69% vs. HO 50% p<0.05). HY n=12, HO n=12, FO n=12. Analyzed by Two way ANOVA.

### PPARγ agonist improves phagocytosis and receptor expression

Given the pronounced age and frailty related reduction in important scavenger receptors CD14 and CD36 we investigated the use of PPARγ agonist rosiglitazone as a potential intervention to rescue phagocytic function in MDM from frail older adults. Pre-treatment with rosiglitazone for 16 hours significantly increased both the percentage of MDM expressing CD36 (26% vs. 40%, p=0.0017, Figure 4a) and CD14 (52% vs. 68% p=0.0068, Figure 4c) but also increased the MFI of expression of both receptors (p=0. 0156 and p=0.0455, fig 4b, c). Rosiglitazone pre-treatment also significantly improved phagocytosis of both *H. influenzae* (38% vs. 50%, p=0.0045, Figure 4e) and *S. pneumoniae* (70% vs. 85%, p=0.0156, Figure 4g) in frail older MDM, expressed both as percentage of MDM phagocytosis and MFI levels (p=0.0156 and p=0.0162 Figure 4f, h).

**Figure 4.**
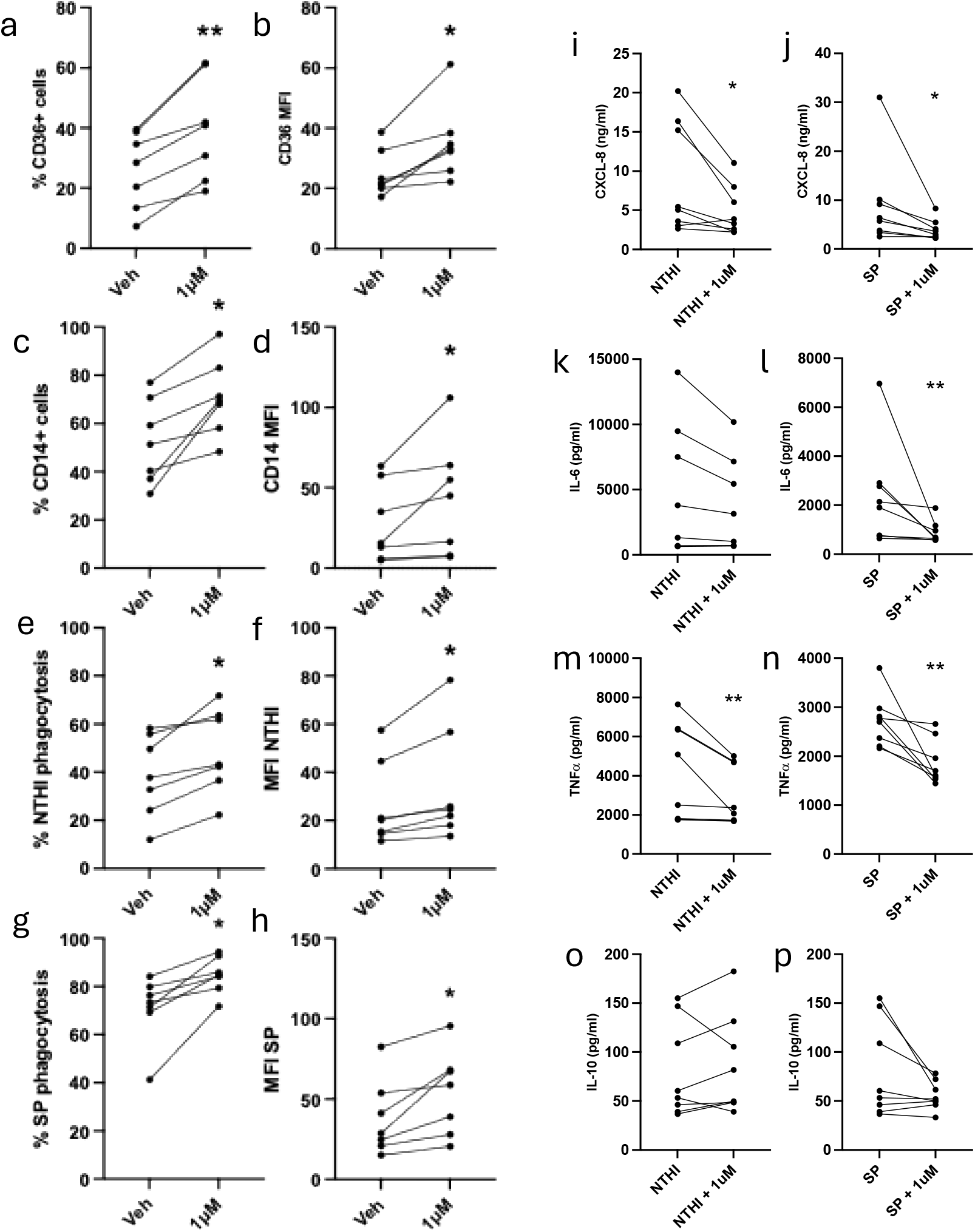
Effect of rosiglitazone on macrophage function. MDM from frail older adults were treated with 1uM rosiglitazone (or vehicle control 0.01% DMSO) for 16 hours, then function measured. Rosiglitazone induced a significant increase in (A) % MDM expressing CD36 (p<0.05), (B) CD36 MFI (p<0.05), (C) % MDM expressing CD14 (p<0.05), (D) CD14 expression (p<0.05), (E) % NTHI phagocytosis (p<0.05), (F) MFI of NTHI phagocytosis (p<0.05), (G) % SP phagocytosis (p<0.05), (H) MFI of SP phagocytosis (p<0.05). Rosaglitazone inhibited the release of (I) CXCL-8 (p<0.05), (K) IL-6 (p<0.05), (M) TNFa (p<0.01), but not (O) IL-10 (p>0.05) induced by NTHI. Rosaglitasone also inhibited the release of (J) CXCL-8 (p<0.05), (L) IL-6 (p<0.01), (N) TNFa (p<0.01) but not (P) IL-10 (p>0.06) induced by SP. n=7 Analysed by Wilcoxon test.

Cytokine release after rosiglitazone treatment in the presence of *H. influenzae* resulted in a significant decrease in the release of CXCL-8 (p=0.0234, Figure 4i) and TNFα (p=0.0078, Figure 4m), and in the presence of *S. pneumoniae* resulted in significant decreases in the release of CXCL-8 (p=0.0156, Figure 4j), IL-6 (p=0.0078, Figure 4l) and TNFα (p=0.0078, Figure 4n). No significant differences to IL-10 release were identified (Figure 4).

### Throat swab microbiome diversity is reduced in frail older adults

The α-diversity (within sample diversity) of the throat swabs was significantly reduced in frail older adults compared to non-frail older and healthy young adults, as indicated by Shannon diversity (fewer different organisms, Figure 5A). The minimal model demonstrated that the age, GDF, VEGF and CRP variables were significantly related to bacterial community structure, together explaining 15% of the variance in community structure (Fig 5B and Supplementary Figure 10). Clinical frailty score and subject group were tested individually to avoid confounding and explained 5% and 7% of the variance in community structure respectively. The most abundant species in all samples were *Streptococcus* and *Prevotella* species (Fig 5C) as seen in other studies of the healthy respiratory tract microbiome^39^. A *Gemella* and a *Selenomonas* OTU were both significantly reduced in frail older compared to either healthy older or healthy younger adults (Supplementary Figure 10B and C). Comparing non-frail older with frail older adults, two more OTUs were significantly reduced in frail older adults, *Segatella* and *Lautropia (*Supplementary Figure 10d, e). Indicator species analysis confirmed these differences and showed that in frail older adults, 97 OTUs were absent compared to healthy young samples, and in non-frail older adults 4 OTUs were absent. A *Streptococcus* and a *Prevotella* OTU were found only in frail older adult samples (Fig 5C). None of these named OTUs correlated negatively with sample biomass and are therefore expected not to be DNA kit contaminants^40^.

**Figure 5.**
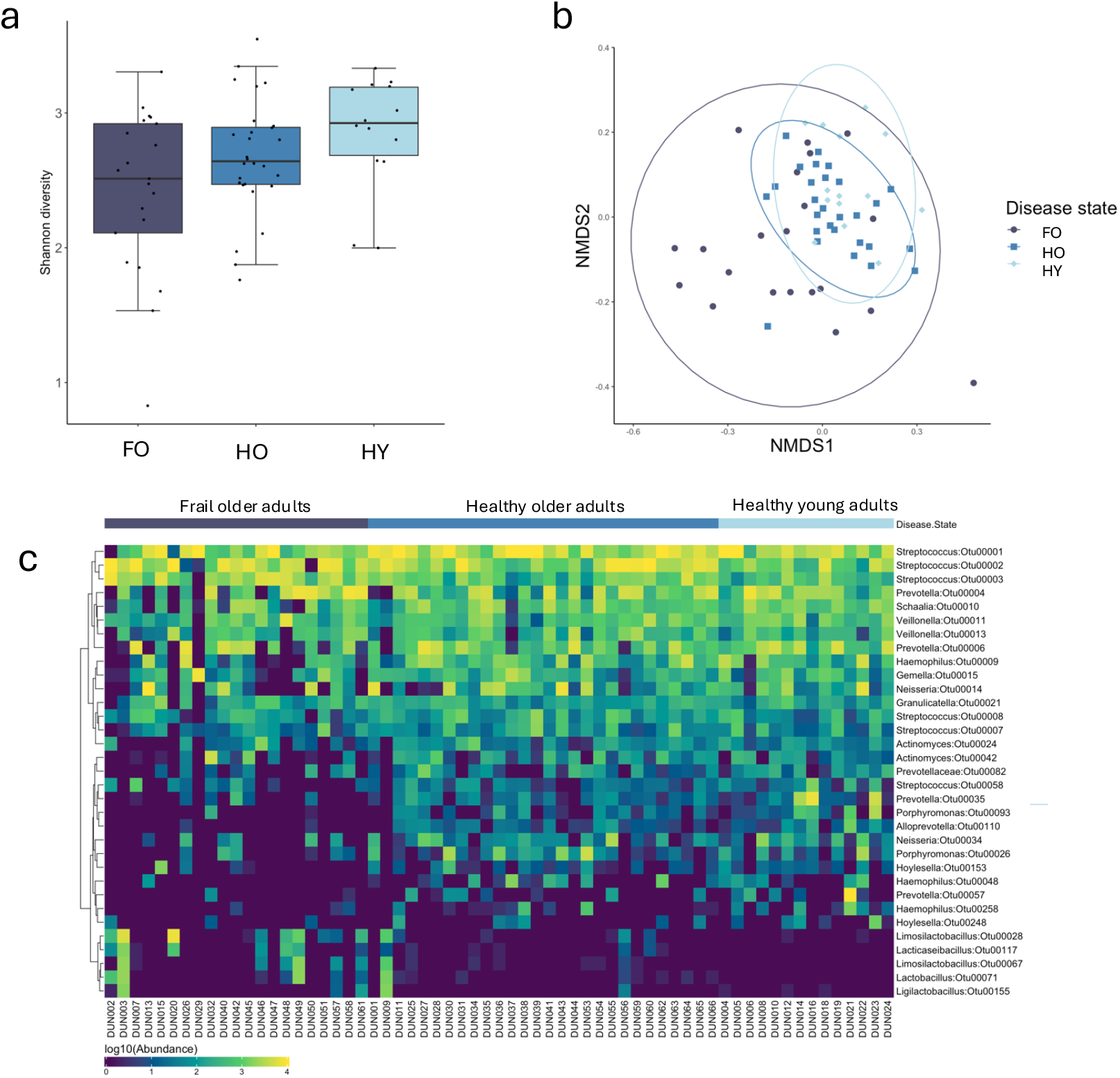
Oropharynx microbiome. Swabs from the oropharynx of participants underwent 16s rRNA gene sequencing for microbiome analysis. A shows Shannon diversity, indicating reduced alpha diversity in frail older adults (p=0.04716, Kruskall wallis chi-squared test). B shows beta diversity. C shows abundance of bacterial species (y axis) against sample type (x axis) grouped by participant type along the top. Indicates loss of diversity in frail older adults.

## Discussion

Frailty is recognized as a state increased vulnerability to infection, with evidence increasing that this is shaped by reduced immune competence. In this study we examined macrophage effector function alongside oropharyngeal microbial communities in healthy young, healthy older and frail older adults. Our study reveals that MDM from frail older adults exhibited reduced phagocytosis of relevant respiratory pathobionts and reduced efferocytosis of apoptotic neutrophils, accompanied by exaggerated release of pro-inflammatory cytokines. In parallel, oropharyngeal samples from frail adults displayed reduced bacterial diversity and altered community compositions. Whilst these results do not establish causality, the co-existence of immune dysfunction and microbial dysbiosis highlights a coupled but unresolved shift at the interface between host and microbe, which leads to increased susceptibility to infection.

To date, our understanding of macrophage ageing is from murine populations, with the changes occurring in human tissue and blood derived cells unclear ^13^. The data we present here extends our knowledge of macrophage inflammageing in humans, with the addition of a clinically frail population with the increased risk of infection and disease in this group, compared to chronological ageing alone. Impairment of macrophage function has been implicated in the pathogenesis of many diseases, suggesting it is key for maintaining health as we age. Our study identifies both age and frailty related declines in macrophage phagocytosis and efferocytosis that is not explained by altered mitochondrial activity. Instead, these changes were linked to reduced expression of key scavenger receptors CD36 and CD14, and over-production of pro-inflammatory cytokines, reinforcing the idea that macrophage dysfunction contributes to the heightened inflammatory state typical in frailty.

Our older adult population was non-frail but still displayed comorbidities typical of the over 65 population, including hypertension and hypercholesteremia. This group displayed increased systemic GDF-15 compared to young adults, a characteristic of normal ageing ^41^, but other systemic cytokines remained low, including CRP. Our frail older adult population had an average CFS score of 6 (moderate frailty) and had higher incidence of comorbidities including cardiovascular disease, arthritis, diabetes and hypertension, characteristic of a frail population. This group displayed increased systemic GDF-15, VEGF, CRP and TIMP-1 compared to non-frail older adults, and young adults, indicative of higher levels of inflammageing linked to frailty^42^. Our young adult population displayed no comorbidities and had low circulating cytokine levels.

We identified distinct features of the upper respiratory microbiome in our frail adult population, including reduced α diversity, consistent with a loss of bacterial richness. These community changes were accompanied by higher levels of pro-inflammatory cytokines typically elevated in frailty. This indicates parallel alterations in both microbial and host immune profiles associated with age and frailty. The interplay between the commensal microbiome and innate immunity is important for understanding ageing. The microbiome plays critical roles in training the immune system, while the immune system itself maintains a healthy microbiome ^43^. Indeed, changes in the lung microbiome with advanced age is linked to lung function decline ^44^. Lower diversity is associated with less colonisation resistance and easier establishment of invasive pathogens, together with a pro-inflammatory environment these observations may start to explain the higher infection risk (both incidence and morbidity) in both healthy older and frail populations. We have identified microbial biomarkers of frailty and healthy ageing in the respiratory tract microbiome alongside a loss of diversity in the respiratory microbiome.

Impaired macrophage phagocytosis in older adults appears exacerbated in frail older adults, alongside impaired efferocytosis and elevated pro-inflammatory cytokine release. Impaired macrophage phagocytosis has implications that range from increased susceptibility to infection, to impaired inflammatory responses. Indeed, impaired phagocytosis is linked to acute respiratory disease ^45^, chronic respiratory disease ^35, 46^, autoimmune disease ^47^ and other conditions. Efferocytosis is also key for the resolution of infection and inflammation, removing potentially necrotic cells from causing further tissue damage, and inducing an anti-inflammatory tissue environment ^48^. Impaired efferocytosis or even slowing of this process can promote inflammation due to elevated pro-inflammatory cytokine levels, and DAMP release from dying cells. Our frail older adult MDM consistently show increased pro-inflammatory cytokine release, both at baseline, and after phagocytosis and efferocytosis – confirming this inflammatory profile seen in these cells. Efferocytosis also failed to stimulate release of IL-10 from frail older adult MDM – a key cytokine in resolution of inflammation ^36^. This likely contributes to increased levels of inflammation in the frail population, indicative of the condition.

Our data supports that of murine studies, where impaired phagocytosis ^11, 12^ and efferocytosis ^49^ results in chronic inflammation. In mice, phagocytosis positively corelates with increased murine lifespan ^50^, implicating impaired phagocytosis in the onset of life-limiting conditions. Our data extends human studies showing impaired MDM phagocytosis of inert beads by older adults ^51^ and impaired bacterial killing of *S. pneumoniae* by frail older adult MDM dependent on PI3K-AKT signaling ^52^. Our data further cements impaired macrophage function as a driver of increased infection risk in aged and frail adults.

To determine a mechanism behind defective macrophage function in frail older adults, we concurrently assessed MDM expression of phagocytic markers at baseline, and after phagocytosis. We found that both older adult MDM and frail older adult MDM had reduced CD36 expression at baseline and after phagocytosis of *H. influenzae*, and reduced expression of CD14 despite phagocytic prey. CD36 is a key efferocytic receptor ^38^ that can also form dimers with TLR2 and TLR4 to act as a phagocytic receptors and stimulate pro-inflammatory cytokine release ^53^. Reduced expression in older and frail older adult MDM may be implicated in their reduced ability to phagocytose/efferocytose. CD14 is a receptor for LPS on bacterial cell walls, but can also bind apoptotic cells and other ligands ^37^. Impaired scavenger receptor expression, and post-phagocytic responses by MDM implicates multiple intracellular mechanisms by which these receptors are produced, trafficked, and recycled within the cell. CD36 activation by apoptotic cells induces activation of PPARγ, a transcription factor involved in lipid metabolism, which in turn upregulates CD36 expression by the cell, in order to participate in further phagocytosis ^54^. In mice, instillation of apoptotic cells after bleomycin injury results in increased PPARγ in AM, resulting in increased CD36 expression which were inhibited by PPARγ antagonists ^55^. PPAR deficient murine macrophages display increased pro-inflammatory cytokine secretion, reduced IL-10 and have impaired phagocytosis compared to WT cells, implicating PPARγ as key in macrophage function ^56^.

We therefore attempted pharmacological rescue experiments to provide additional mechanistic insight. Rosiglitazone, a PPARv agonist, restored phagocytic function against both of both *H. influenzae* and *S*.*pneumoniae*, reduced pro-inflammatory cytokine release and elevated both CD36 and CD14 expression by frail older adult MDM. These effects are consistent with a role for PPARγ in regulating lipid metabolism and receptor recycling These results highlight a tractable pathway for intervention and a promising pathway for further exploration in age-related immune decline. Whilst Rosiglitazone is no longer widely used in clinical practice due to off-target effects, similar compounds can manipulate this pathway. Piaglitazone can promote an anti-inflammatory phenotype of mouse bone marrow derived macrophages via PPARγ activation ^57^ supporting this idea. Knockdown of MYC in human MDM, a gene that upregulates PPARγ simulates this decline in macrophage phagocytosis ^58^ confirming our theories.

Several limitations should be acknowledged. Oropharyngeal swabs cannot directly assay the lower respiratory tract, though they represent an accessible niche relevant to respiratory risk with several studies supporting their clinical significance without direct sampling of the lung^59^. Recruitment of frail older adults from the hospital environment introduces potential confounding factors; comorbidities and acute illness. Medication use such as proton pump inhibitors, known to be a confounding factor in gastrointestinal microbiome studies, could not be controlled for in this study. The use of monocyte-derived macrophages over alveolar macrophages is a limitation, however due to the vulnerability of frail populations this experiment is not feasible on humans. However we have previously shown that MDM function correlates with AM function ^35^, lending importance to this data regardless. Finally, this study is cross-sectional by nature, limiting our power to draw causal inference directly between immune and microbiome alterations.

As such, our study gives good justification for longitudinal study in relevant/ at risk populations in frail and healthy older people of community acquired pneumonias, with a focus on disentangling cause and effect of the immune system and microbiome. Population studies of PPARγ inhibitors as an intervention for restoration of a younger microbiome composition and in preventing respiratory tract infections.

Overall, we show a systemic defect in monocyte-derived macrophage function in older adults, which is enhanced in frailty. Frailty was also associated with reduced bacterial diversity in the oropharynx. The relationship between macrophage function and bacterial richness remains to be determined, however targeting impaired immune cell function, such as with PPARγ agonists may improve responses to infection and thus reduce infection risk in frail older adults.

### Recourse Availability

Further information and requests for resources should be directed to and will be fulfilled by the lead contact, Kylie Belchamber. 16S rRNA gene sequence data has been submitted to the European Nucleotide Archive under bioproject ID PRJEB107714

## Acknowledgements

Authors would like to acknowledge the Birmingham Inflammation Research Facility for assisting in patient appointments. This study was funded by the Vivensa Foundation AIS2210-7.

## Author contributions

KBRB, ES, TJ, DP, MJC and AS conceptualized and designed studies. KBRB, KPY, OST, FSG recruited participants. KBRB performed experiments. KBRB, SP, MJC analysed data. KBRB wrote the paper. All authors read and edited the paper.

## Supplemental information

Document S1

Supplementary figures S1-10

---

**Supplementary Figure 1.**
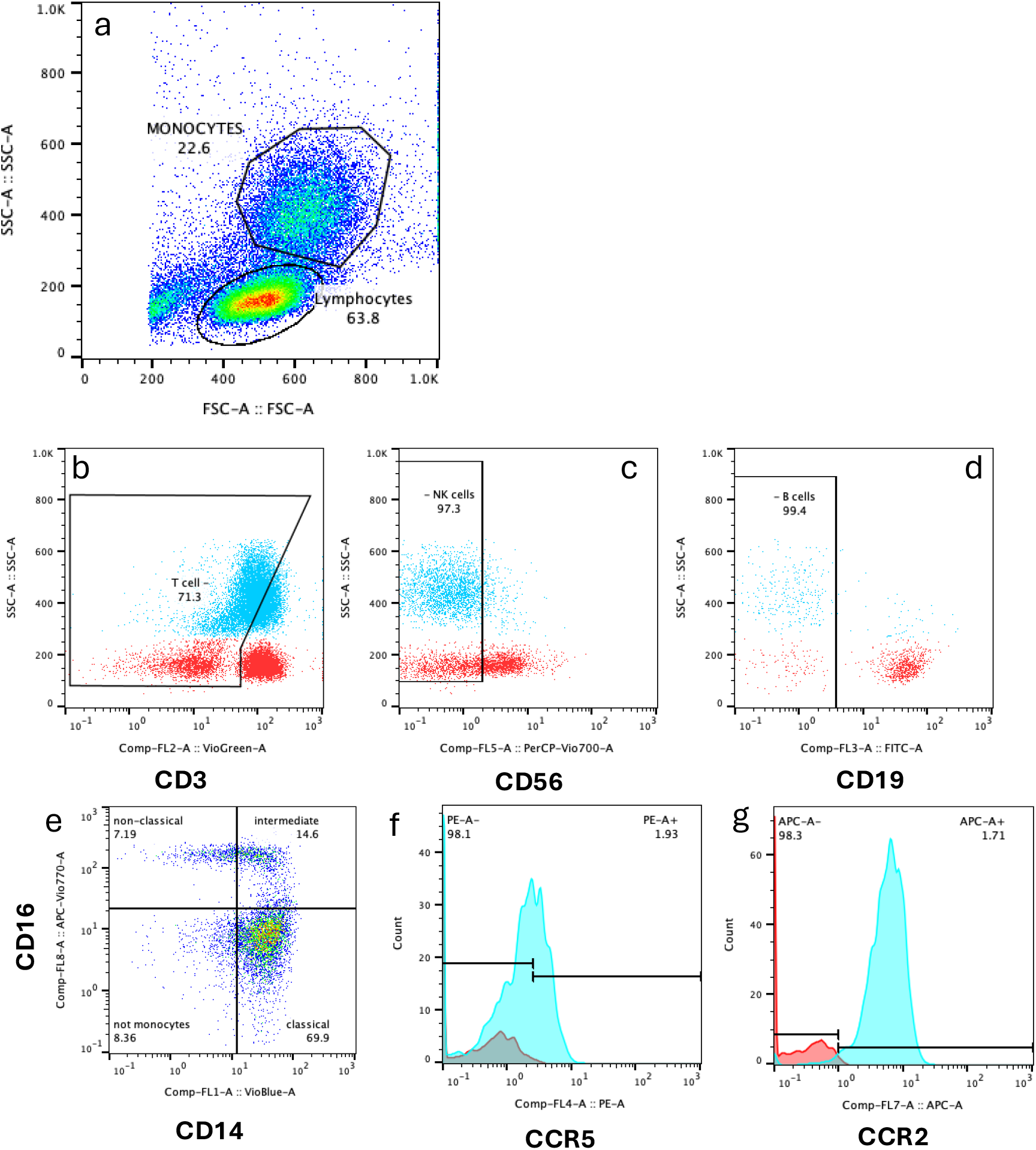
Monocyte gating strategy. Whole blood was assessed for monocyte populations by flow cytometry. a) FSC vs SSC gating to identify monocyte (and lymphocyte) populations. b) Monocyte population (blue) excluding CD3+ T cells (red shows lymphocyte population). c) CD3 - monocyte population excluding CD56+ NK cell population. d) Monocyte population excluding CD19+ B cell population. e) CD14+ CD16+ monocyte populations, separated based on monocyte phenotype. f) CCR5 expression by monocytes (blue) compared to –CCR5 FMO control (red). g) CCR2 expression by monocytes (blue) compared to –CCR2 FMO control (red).

**Supplementary Figure 2.**
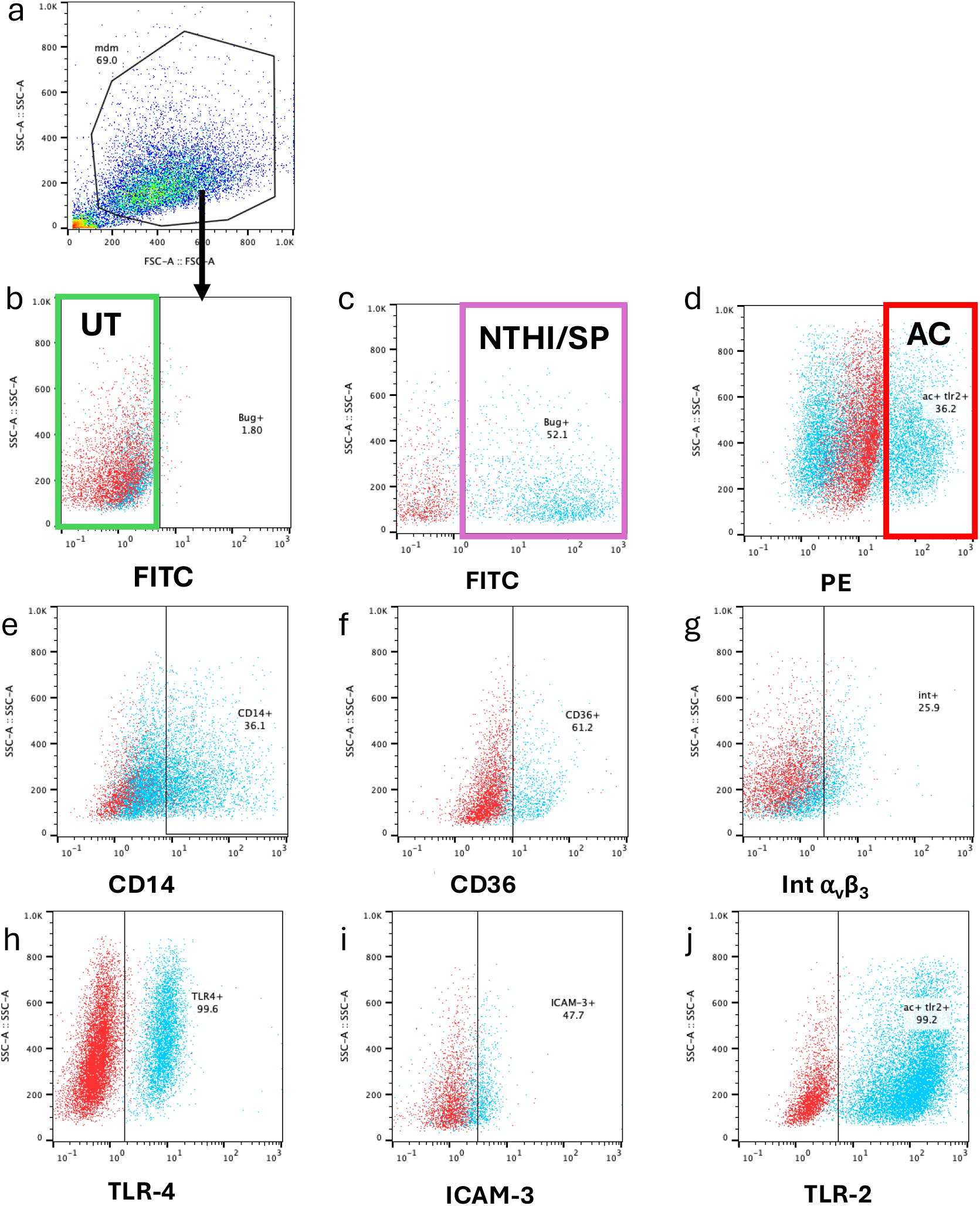
MDM gating strategy. MDM were assessed for phagocytosis or efferocytosis, and scavenger receptor expression by flow cytometry. a) FSC vs SSC of MDM population. b) Untreated (UT) macrophages were assessed in the FITC-gate, where blue indicates UT control, and red indicates FMO control c) NTHI or SP+ macrophages were assessed in the FITC+ gate (blue=sample, red=unstained) d) Efferocytic macrophages were assessed in the PE+ gate. Expression of scavenger receptors were assessed in these assigned gates, were blue indicates a stained sample, and red indivates FMO control e) CD14 expression. f) CD36 expression. g) Integrin α_v_β_3_ expression. h) TLR4 expression. i)ICAM-3 expression. j) TLR-2 expression (not assessed for efferocytosis, as both assessed on PE channel.

**Supplementary Figure 3.**
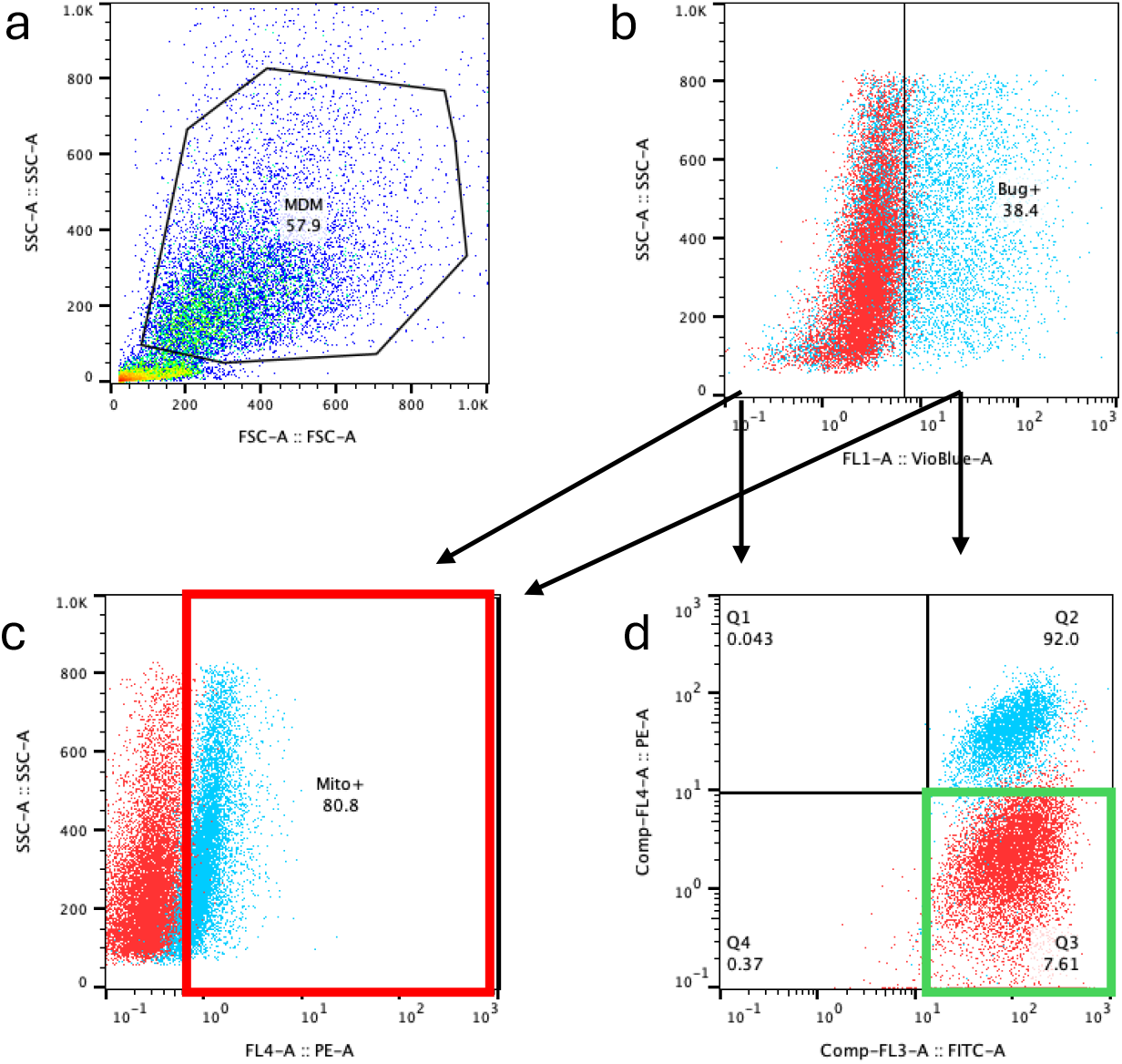
Mitochondrial function gating strategy. MDM were assessed for mitochondrial function by flow cytometry. a) FSC vs SSC of MDM population. b) Untreated MDM (red) were assessed for mitosox and JC-1 in the Vioblue-channel. Phagocytic MDM (NTHI or SP+) were assessed in the Vioblue+ channel. c) Samples were assessed for mitosox expression by MFI, in the PE+ gate. Red shows unstained control, and blue shows a stained sample. d) JC-1 was used to assess mitochondrial membrane potential, expressed as %green monomers. JC-1 fluoresces in the PE channel as red aggregates within mitochondria, indicated polarized mitochondria. JC-1 fluoresces in the FITC channel as green monomers in depolarized mitochondria. Red shows CCCP positive control sample, and blue shows un-stimulated control.

**Supplementary figure 4.**
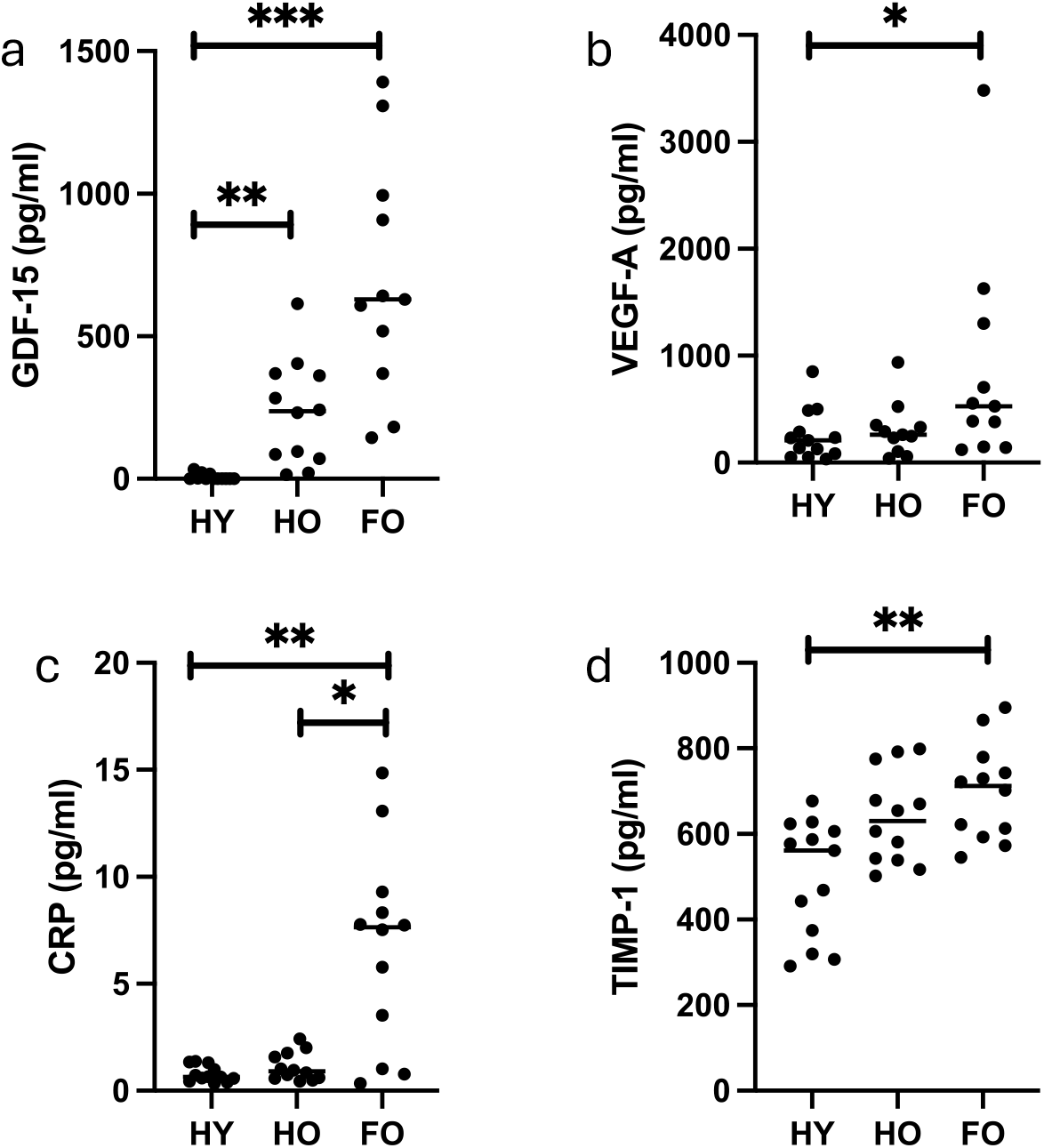
Serum cytokines across patient populations. (A) Compared to HY participants, GDF-15 was significantly increased in both HO (6±3pg/ml vs 233±53pg/ml, p<0.01) and FO participants (700±125pg/ml, p<0.001). (B) Compared to HY participants, VEGF-A was significantly increased in FO participants (254±65pg/ml vs. 853±300pg/ml, p<0.05). (C) Compared to HY and HO participants, CRP was significantly increased in FO participants (0.78±0.1pg/ml HY vs. 1.1±0.2pg/ml HO vs. 6.7±1.3pg/ml FO, p<0.01 and p<0.05 respectively). (D) Compared to HY participants, TIMP-1 was significantly increased in FO participants (498±38pg/ml vs. 699±33pg/ml, p<0.001). HY n=12, HO n=12, FO n=12. Analysed by Kruskall-Wallis test.

**Supplementary Figure 5.**
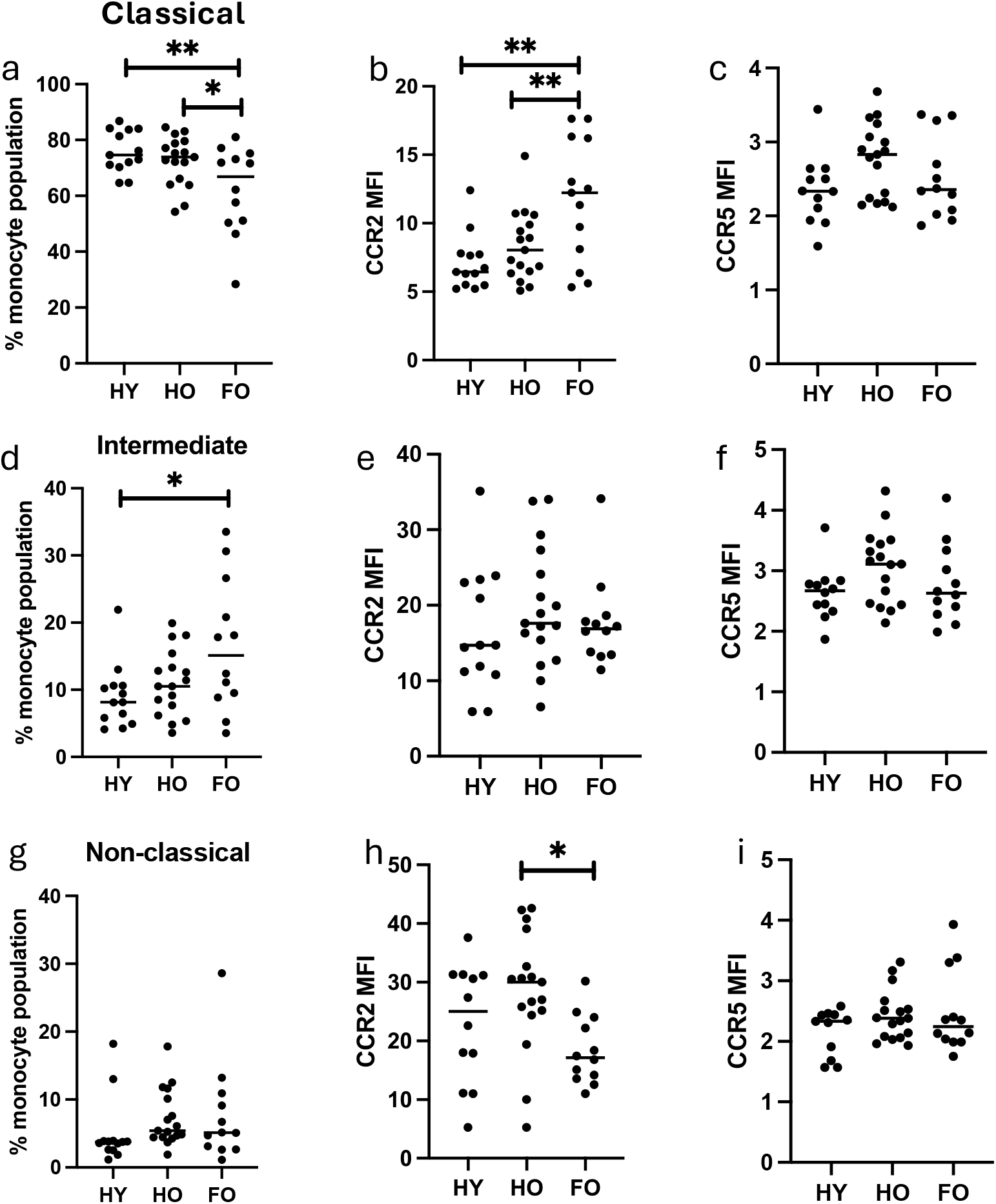
Monocyte populations and activation. (A) The percentage of CD14^+^ CD16^−^ classical monocytes was significantly decreased in FO adults compared to both HY and HO adults (HY 76±2% vs HO 72±2% vs FO 62±5%, p<0.01 and p<0.05). (B) Expression of CCR2 on classical monocytes was significantly increased in FO adults (HY 7.1±0.57 vs HO 8.4±0.6 vs FO 11.7±1.2 MFI, p<0.01 and p<0.01. (C) Expression of CCR5 was not different on classical monocytes. (D) The percentage of CD14^++^ CD16^+^ intermediate monocytes was significantly increased in FO adults compared to HY adults (HY 9.0±1.3% vs HO 11.0±1.2% vs FO 16.5±2.8%, p<0.05). (E) Expression of CCR2 did not change between patient groups. (F) Expression of CCR5 did not differ between patient groups. (G) The percentage of CD16^+^ CD14^−^ non-classical monocytes did not differ between groups (p>0.05). (H) Expression of CCR2 was significantly decreased in FO adults compared to HO adults (HY 22.9±3 vs HO 28.4±2.5 vs FO 18.4±1.7 MFI, p<0.05). (I) Expression of CCR5 did not differ between groups. HY n=12, HO n=17, FO n=12. Analysed by One Way ANOVA.

**Supplementary Figure 6.**
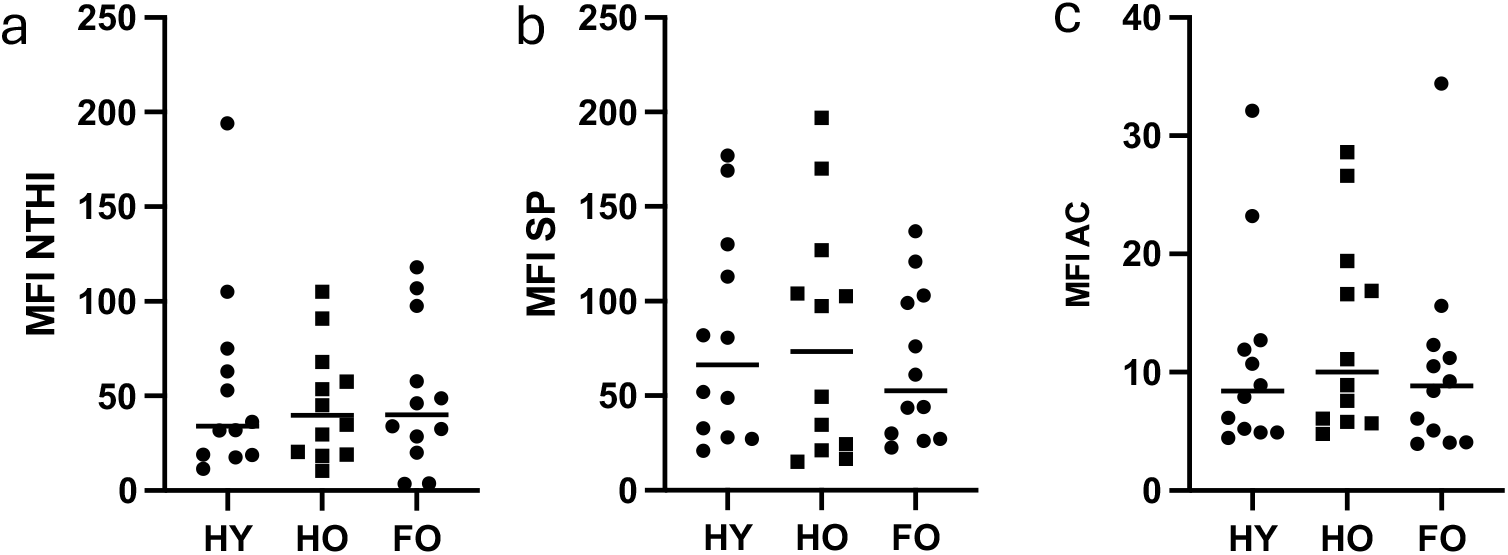
MFI of phagocytosis. The amount of prey phagocytosed expressed as MFI did not differ between subject groups for (A) NTHI (p>0.05), (B) SP (p>0.05) or (C) efferocytosis (p>0.05).

**Supplementary Figure 7.**
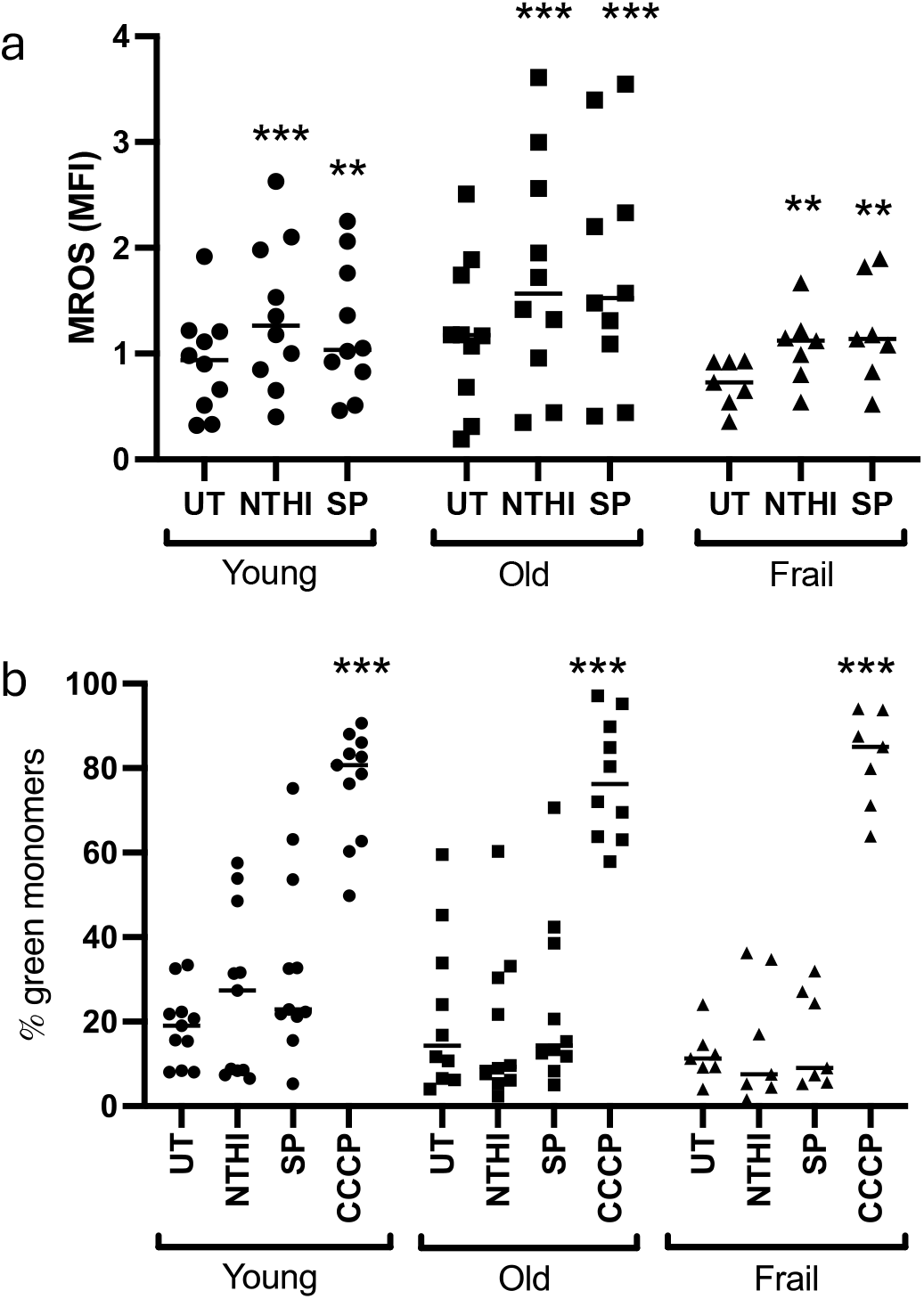
Mitochondrial health. (A) Across all patient groups, phagocytosis of NTHI significantly increased mROS compared to UT cells (p<0.01-0.001). There were no significant differences between patient groups. (B) MMP measured by JC-1 assay and presented as % green monomers. The positive control CCCP significantly increased % green monomers in all subject groups (p<0.001), however there was no change induced by phagocytosis of either NTHI or SP (p>0.05). There was no difference between patient groups (p>0.05). HY n=11, HO n=11, FO n=7. Analysed by 2 Way ANOVA.

**Supplementary figure 8.**
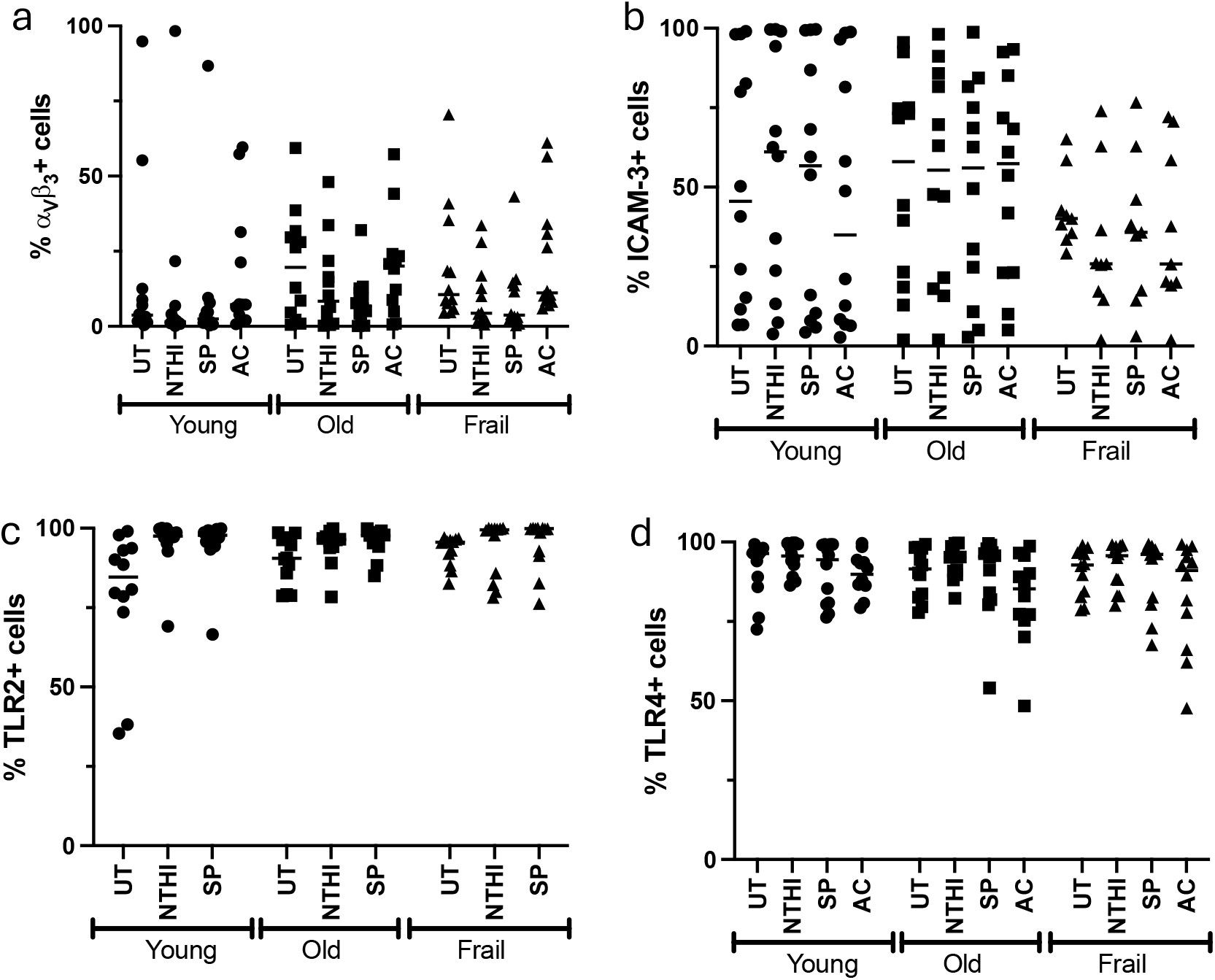
% expression of receptors. Expression of (a) integrin α_v_β_3_, (b) ICAM-3, (c) TLR2 and (d) TLR4 at baseline (UT) and after phagocytosis/efferocytosis by MDM. There was no significant change in receptor expression after phagocytosis, or between patient groups. N=12 for all groups. Analysed by 2 Way ANOVA.

**Supplementary figure 9.**
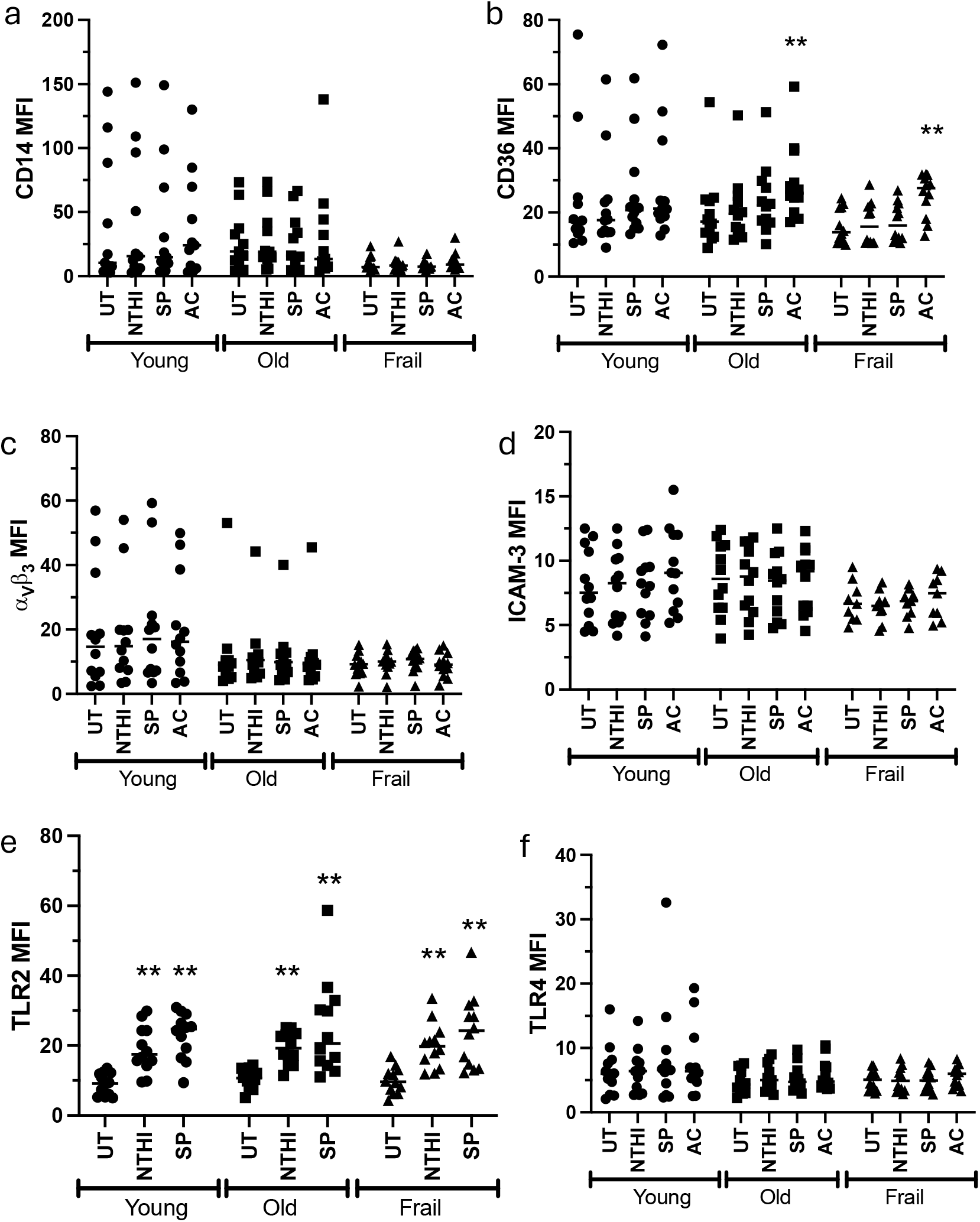
MFI of receptors. MFI of (a) CD14, (b) CD36, (c) integrin α_v_β_3_, (d) ICAM-3, (e) TLR2 and (f) TLR4 at baseline (UT) and after phagocytosis/efferocytosis by MDM. There was no significant change in receptor expression after phagocytosis, or between patient groups. N=12 for all groups. Analysed by 2 Way ANOVA.

**Supplementary figure 10.**
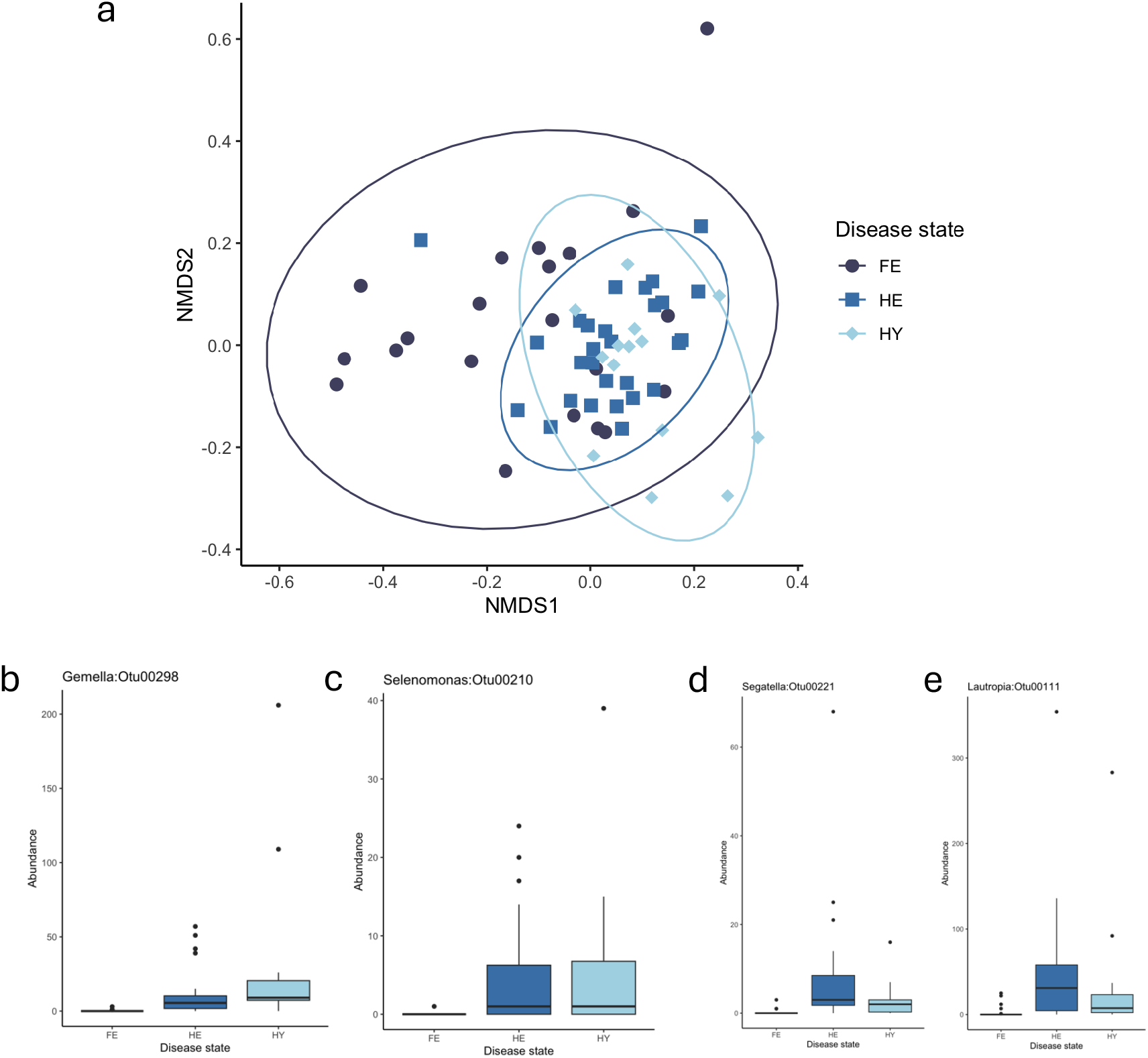
Beta diversity and differently abundant OTUs. A – Non-metric multidimensional scaling (NMDS) ordination plot. Each point is a sample and distance between samples represents the dissimilarity between microbial communities within each sample (beta diversity). Ellipses were fitted using envfit in vegan. Disease state accounts for 41% of the variation in beta diversity (p = 0.013) B - OTU plot for *Gemella* indicating it is less abundant in frail older adults. C shows OTU plot for *Selenomonas* indicating it is less abundant in frail older adults. D shows OTU plot for *Segatella*. Indicating it is more abundant in healthy older adults. E shows OTU plot for *Lautropia* indicating it is more abundant in healthy older adults.

