## Supplementary text for "Macrophage Phagocytic Impairment is Associated with Dysbiosis of the Respiratory Microbiome in Frail Older Adults"

### Sequence processing

Demultiplexed FASTQ sequencing files without primers were processed through mothur version 1.48.2 [1] using the standard operating procedure described in Kozich *et al.* [2]. The files were paired into contigs, and sequences between 420 and 430 nucleotides in length with no ambiguous nucleotides were extracted. These sequences were aligned to the SILVA reference database release 138.2 [3] which was trimmed to only contain the region being sequenced as described in Schloss [4]. This trimmed region was between positions 1 and 17017 of the original database. After aligning and a second length screen, sequences were pre-clustered and chimeras were removed. Remaining sequences were classified into taxa using the Ribosomal Data Project classifier version 19 [5]. Processed sequences were clustered into operational taxonomic units (OTUs) with a 97% similarity threshold. Resulting OTUs were classified and their abundance in each sample counted, producing taxonomy and abundance tables ready for analysis.

All work in mothur was carried out using version 1.48.2 installed on BlueBEAR, the high-performance computing service at University of Birmingham. Each of the 54 central processing units requested were allocated 364.5GB of random access memory.

### Analysis in R

*OTU abundance and taxonomy tables were imported into R version 4.4.2 (R Core Team, 2024) using RStudio version 2024.12.0+467. They were imported as a phyloseq object using the phyloseq package[6]. For the sample metadata of this phyloseq object, a text file was imported to R containing details of DNA extraction, the subject's frailty grouping, age, clinical frailty score, and cytokine levels, among other things.*

To find which OTUs were likely contaminants, the decontam package version 1.26 [7] was used in R. The isContaminant() function with the 'frequency' setting identified OTUs whose abundance was negatively correlated with the concentration of DNA extracted from each sample. These OTUs were removed from all samples.

*To study differences in alpha diversity between frail elder, healthy elder, and healthy young people, decontaminated samples were rarefied to 33000 sequences each. This corresponded to the lowest number of sequences in a sample rounded down to the nearest thousand. Species richness was calculated manually while Shannon and Simpson diversity indices were calculated using the vegan package [8].*

Beta diversity was analysed in four ways using the vegan, phyloseq and ComplexHeatmap packages on the decontaminated rarefied dataset [6, 8, 9]. Firstly, permutational analyses of variance were conducted using the adonis2() function to see which metadata variables, affected community composition. Significant metadata variables were included as adonis2() strata in the main permutational analysis of variance – how microbiome composition varies with frailty. Two measures of frailty were used: either frail elder, healthy elder, and healthy young groups, or a clinical frailty score from 1-8. Then, an annotated heatmap was made to see whether OTUs with at least 5% abundance in at least one sample could distinguish between the three groups of people. The 5% abundance threshold was chosen following Socransky [10]. A stacked barplot was also created to assess the abundance of different taxonomic classes between samples. Finally, non-metric multidimensional scaling was carried out using Jaccard distance to measure microbiome dissimilarity.

Differentially abundant OTUs were identified using ANCOM-BC2 and DESeq2 [11, 12]. While DESeq2 has a high true positivity rate, ANCOM-BC2 has a much lower false discovery rate and corrects for the zero-inflated nature of microbiome data [13, 14]. To make DESeq2 more applicable to microbiome abundances, the OTU table was normalised by geometric mean of pairwise ratios before using DESeq2 – as proposed by Chen *et al.* (2018) and implemented in the GUniFrac package[15]. In addition, indicator species analysis was performed using the indicspecies package [16] to determine which OTUs are more likely to be found in frail people than healthy people.

Before processing: 65100 ± 8490 sequences per sample
After processing: 50533 ± 9341 sequences per sample
